# Neuronal autophagy regulates presynaptic neurotransmission by controlling the axonal endoplasmic reticulum

**DOI:** 10.1101/2020.07.06.189522

**Authors:** Marijn Kuijpers, Gaga Kochlamazashvili, Alexander Stumpf, Dmytro Puchkov, Max Thomas Lucht, Eberhard Krause, Dietmar Schmitz, Volker Haucke

**Affiliations:** Leibniz-Forschungsinstitut für Molekulare Pharmakologie (FMP), 13125 Berlin, Germany; Freie Universität Berlin, Faculty of Biology, Chemistry, Pharmacy, 14195 Berlin, Germany; Charité Universitätsmedizin Berlin, corporate member of Freie Universita□t Berlin, Humboldt-Universita□t zu Berlin, and Berlin Institute of Health, 10117, Berlin, Germany

## Abstract

Information processing in the brain is encoded as electrical impulses in neurons that are relayed from the presynaptic compartment to postsynaptic neurons by regulated neurotransmitter release. Neurons are known to rely on autophagy for the removal of defective proteins or organelles to maintain synaptic neurotransmission and to counteract neurodegeneration. In spite of its importance for neuronal health, the physiological substrates of neuronal autophagy in the absence of proteotoxic challenge have remained largely elusive. We use knockout mice conditionally lacking the essential autophagy protein ATG5 and quantitative proteomics to demonstrate that loss of neuronal autophagy causes the selective accumulation of tubular endoplasmic reticulum (ER) in axons, resulting in increased excitatory neurotransmission and compromised postnatal viability *in vivo*. The gain in excitatory neurotransmission is shown to be a consequence of elevated calcium release from ER stores via ryanodine receptors accumulated in axons and at presynaptic sites. We propose a model in which neuronal autophagy controls axonal ER calcium stores to regulate neurotransmission in healthy neurons and in the brain.

## INTRODUCTION

Information processing in the brain critically relies on the relay of information from a presynaptic neuron to the postsynapse via regulated neurotransmitter release. This process is triggered by the action potential (AP)-triggered calcium-driven exocytic fusion of neurotransmitter-containing synaptic vesicles (SVs) at active zone (AZ) release sites (Jahn and Fasshauer, 2012; Sudhof, 2013). Exocytic SV fusion is followed by endocytosis of SV membranes and the reformation of functional SVs to replenish the SV pool (Haucke et al., 2011; Murthy and De Camilli, 2003; Rizzoli, 2014). The efficacy of neurotransmitter release among other mechanisms is modulated by presynaptic calcium influx via voltage-sensitive calcium channels located at AZs, calcium efflux and sequestration (Nanou and Catterall, 2018; Neher and Sakaba, 2008), as well as calcium-induced calcium release from internal endoplasmic reticulum (ER) stores located in the axon and at presynaptic sites (Bezprozvanny and Kavalali, 2020; Galante and Marty, 2003; Irie and Trussell, 2017).

As neurons are long-lived postmitotic cells, the majority of their synapses need to be maintained for the entire lifespan of the organism (Cajigas et al., 2010). To prevent neuronal and synaptic dysfunction neurons have evolved mechanisms for the removal of toxic or defective proteins and organelles to maintain regulated neurotransmission and the integrity of their functional proteome. Among these mechanisms are the degradation of cytoplasmic proteins via the ubiquitin-proteasome system (Hakim et al., 2016), the lysosomal turnover of membrane proteins, and macroautophagy (simply referred to as autophagy hereafter), a cellular process by which defective proteins and organelles are degraded through sequestration within autophagosomes and delivery to lysosomes (Hill and Colon-Ramos, 2020; Nikoletopoulou and Tavernarakis, 2018; Vijayan and Verstreken, 2017). The formation of autophagosomes requires the ATG12-ATG5-ATG16L protein complex that conjugates proteins of the LC3 family to the forming autophagosome membrane (Ariosa and Klionsky, 2016; Hill and Colon-Ramos, 2020; Nikoletopoulou and Tavernarakis, 2018; Vijayan and Verstreken, 2017). In neurons autophagy has been implicated in diverse processes ranging from development including signaling via neurotrophins (Andres-Alonso et al., 2019; Kononenko et al., 2017) to the pathogenesis of major neurodegenerative disorders (Moreau et al., 2014; Nixon, 2013; Ravikumar et al., 2010; Sarkar et al., 2007; Stavoe and Holzbaur, 2019)}(Moreau et al., 2014; Nixon, 2013; Ravikumar et al., 2010). The importance of the autophagy system in the brain is further emphasized by the fact that knockout of core ATG proteins such as ATG5 or ATG7 induces the accumulation of non-degraded protein aggregates, neurodegeneration, and neuronal cell death in mice (Hara et al., 2006; Komatsu et al., 2006; Komatsu et al., 2007). Conversely, induction of autophagy has been suggested to counteract neurodegeneration in disease models (Moreau et al., 2014; Nixon, 2013; Ravikumar et al., 2010; Ravikumar et al., 2004; Williams et al., 2006).

In spite of the general importance of autophagy for neuronal viability and function (Friedman et al., 2012; Hill and Colon-Ramos, 2020; Nikoletopoulou and Tavernarakis, 2018; Vijayan and Verstreken, 2017), the physiological substrates of neuronal autophagy and the mechanisms by which defects in neuronal autophagy affect neuronal and synaptic function are largely unknown. Live imaging studies in cultured neurons and in invertebrate models have established that autophagosomes are formed in distal axons and within the presynaptic compartment (Hill and Colon-Ramos, 2020; Maday and Holzbaur, 2014; Maday et al., 2012). Consistent with this model, presynaptic endocytic proteins such as Endophilin and Synaptojanin have been found to play a role in the induction of neuronal autophagy at presynaptic sites (Azarnia Tehran et al., 2018; Murdoch et al., 2016; Soukup et al., 2016; Soukup and Verstreken, 2017). Distally formed autophagosomes mature during their retrograde axonal transport (Guedes-Dias and Holzbaur, 2019; Stavoe and Holzbaur, 2019) before their fusion with degradative lysosomes enriched in proximal axons and in neuronal somata (Hill and Colon-Ramos, 2020; Maday and Holzbaur, 2014; Maday et al., 2012). In addition to this largely constitutive process of neuronal autophagy (Maday and Holzbaur, 2016), the formation of autophagosomes has been suggested to be facilitated by mitochondrial damage (Ashrafi et al., 2014), neuronal activity (Shehata et al., 2012; Wang et al., 2015), overexpression of aggregation-prone proteins (Corrochano et al., 2012), ROS-induced protein oxidation (Hoffmann et al., 2019), or genetic depletion of key AZ proteins (Okerlund et al., 2017). For example, loss of the AZ protein Bassoon has been shown to trigger the autophagic turnover of SV proteins (Okerlund et al., 2017) and Rab26 has been suggested to target SVs to autophagosomal structures (Binotti et al., 2015), suggesting that SVs and possibly other presynaptic proteins are turned over via neuronal autophagy (Hernandez et al., 2012).

We demonstrate here using knockout mice conditionally lacking the essential autophagy protein ATG5 and quantitative proteomics that loss of neuronal autophagy causes the selective accumulation of tubular endoplasmic reticulum (ER) in axons, resulting in increased excitatory neurotransmission due to elevated calcium release from ER stores via ryanodine receptors. Our findings suggest neuronal autophagy controls axonal ER calcium stores to regulate neurotransmission in healthy neurons and in the brain.

## RESULTS

### Loss of neuronal autophagy in the absence of ATG5 facilitates excitatory neurotransmission and causes premature death *in vivo*

Previously it was demonstrated that early loss of ATG5 in neurons and glial cells throughout the nervous system causes progressive motor deficits and severe neurodegeneration associated with ubiquitin-containing cytoplasmic inclusions (Hara et al., 2006; Komatsu et al., 2006). To determine the physiological consequence of selective ablation of autophagy in neurons in the neocortex and hippocampus, we crossed ATG5*^flox/flox^* mice with a transgenic EMX1-*Cre* line that expresses Cre recombinase in postmitotic excitatory neurons. Conditional ATG5*^flox/flox^*; EMX1-*Cre* KO mice (hereafter termed ATG5-cKO) were born at normal Mendelian ratios (Fig. S1A), but displayed reduced postnatal growth (Figure S1B) and early postnatal lethality between 2 and 6 months of age (Figure 1A). Analysis by immunoblotting revealed a profound loss of ATG5 protein mainly in the cerebral and cerebellar cortex and in the hippocampus that was accompanied by accumulation of the autophagy adaptor and substrate protein p62 (Figure 1B), consistent with prior observations in ATG5*^flox/flox^*; nestin-*Cre* KO mice lacking ATG5 throughout the brain (Hara et al., 2006). Accumulation of p62 in the cortex and hippocampus as well as signs of astrogliosis were observed by confocal imaging in brain slices (Figures 1C; S1C). Moreover, caspase activity was elevated in aged 4 months-old but not in young ATG5-cKO mice (Figure S1D,E). These results show that loss of neuronal autophagy impairs postnatal viability and causes neuronal cell death in mice *in vivo*.

**Figure 1.**
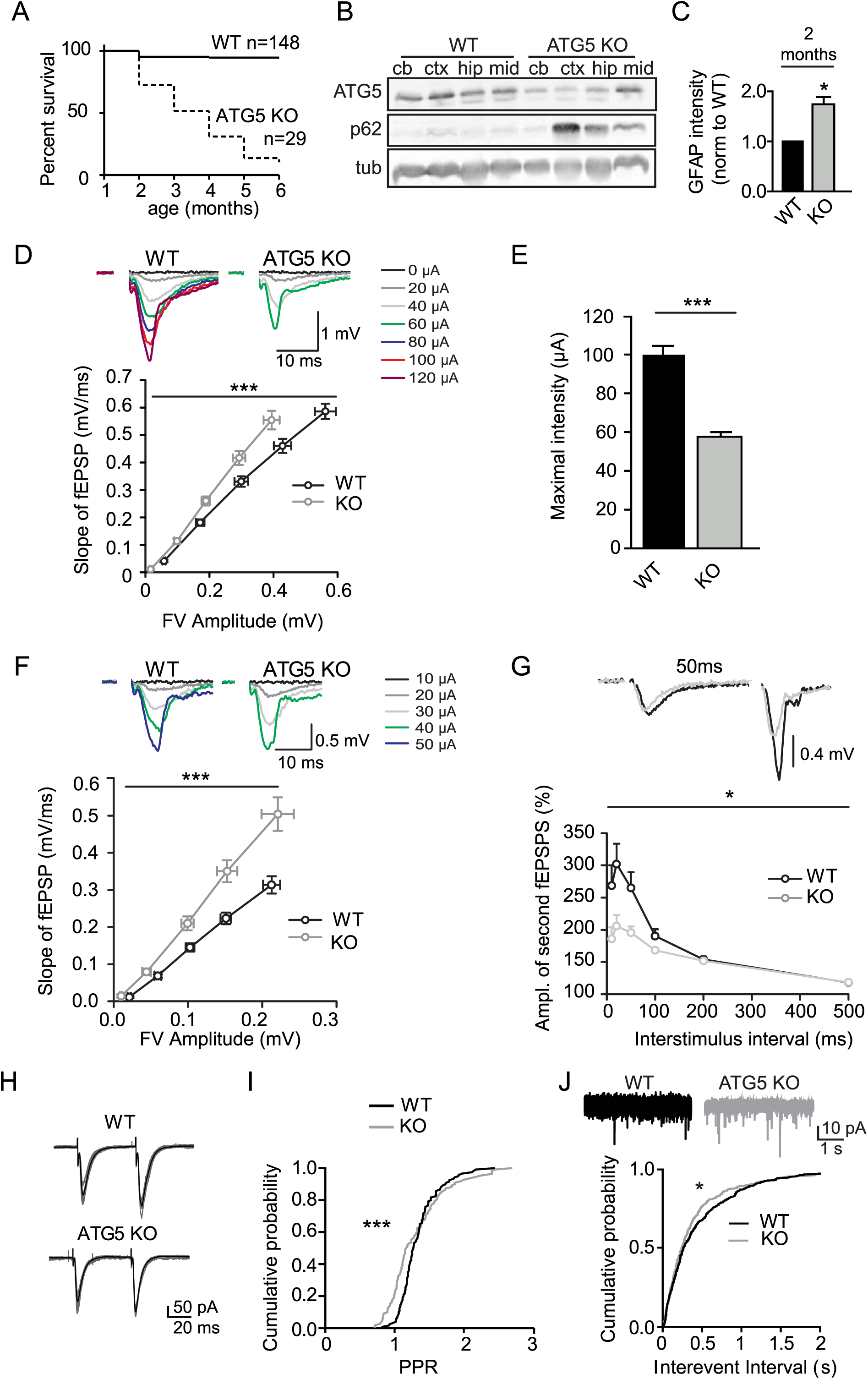
Loss of neuronal autophagy facilitates excitatory neurotransmission. (A) Decreased survival of KO mice conditionally deleted for ATG5 by transgenic expression of Cre recombinase under the telencephalon-specific EMX promoter (ATG5^flox/flox^; EMX1-Cre). (B) Western blot showing ATG5 decrease and p62 accumulation primarily in cortex (ctx) and hippocampus (hip) of ATG5-cKO mice. (C) Quantification of GFAP immunostaining in 6-7 weeks-old control and ATG5-cKO brain slices. Slices taken from 3 mice; one-sample t-test. See also Figure S1C. (D) Basal excitatory neurotransmission measured as relationship between fiber volley (FV) amplitudes and slopes of fEPSPs in WT control (n=24 slices, 12 mice) and ATG5-cKO mice (n=24 slices, 12 mice). Representative fEPSP traces (above) and quantified data are shown. Significant difference between WT control and ATG5-cKO slices encompassing the curve; two-way repeated measures ANOVA. (E) Lower stimulation intensity is required to elicit maximal responses in ATG5-cKO (58.8 ± 2.1 µA) compared to control mice (100.8 ± 4.6 µA); t-test. (F) Basal excitatory neurotransmission measured as relationships between FV amplitudes and slopes of fEPSPs in WT control (n=11 slices, 6 mice) and ATG5-cKO mice (n=10 slices, 6 mice) in the presence of GABA_A_ receptor antagonist picrotoxin (50 µM). Representative fEPSP traces (above) and quantified data are shown. Significant difference between WT control and ATG5-cKO slices encompassing the curve; two-way repeated measures ANOVA. (G) Measurements of paired-pulse facilitation (PPF) in the presence of GABA_A_ receptor antagonist picrotoxin (50 µM) reveal significantly reduced PPF in ATG5-cKO (n=10 slices, 6 mice) compared to control mice (n=11 slices, 6 mice). Representative traces of PPF at 50 ms interstimulus interval (above) and quantified data over a range of interstimulus intervals (10–500 ms), given as ratio of the second to the first response, show reduced facilitation of the second response in ATG5-cKO mice; two-way repeated measures ANOVA. (H) Example traces of voltage clamp recordings with paired extracellular stimulation (50ms ISI) to determine paired pulse ratio (PPR). Gray: individual traces; black: average trace. (I) Cumulative probability shows a more widespread distribution for PPR in ATG5-cKO mice. n=12 (WT) or 11 (KO) slices from 3 animals; Kolmogorov-Smirnov test. (J) Cumulative probability distribution shows decreased interevent intervals for spontaneous EPSCs in ATG5-cKO mice. n=14 (WT) or 17 (KO) slices from 4 animals; Kolmogorov-Smirnov test. All data show mean ± SEM. ns: not significant; p < 0.05; p < 0.001.

To analyze if and how loss of neuronal autophagy in the conditional absence of ATG5 in excitatory neurons affects synaptic transmission, we recorded field excitatory postsynaptic potentials (fEPSPs) of CA3-CA1 synapses in acute hippocampal slices. These measurements revealed elevated basal synaptic transmission in ATG5-cKO mice. The slopes of fEPSPs over fiber volley (FV) amplitudes were significantly increased (Figure 1D) and lower stimulation intensities were required to elicit maximal responses in ATG5-cKO slices (Figure 1E). Elevated basal transmission was not due to altered levels or localization of pre- (i.e. SV proteins) and postsynaptic proteins (Figure S1F,G; and Figs. 2 and 3 below). Moreover, elevated fEPSP slopes over FV amplitudes were also observed in the presence of the GABA_A_ receptor antagonist picrotoxin (Figure 1F), suggesting that elevated excitatory transmission in cATG5-KO slices was not a consequence of impaired synaptic inhibition. We therefore followed the alternative hypothesis that loss of neuronal autophagy facilitates excitatory neurotransmission by increasing presynaptic release probability (Branco and Staras, 2009). Consistently, slices from ATG5-cKO mice showed reduced paired-pulse facilitation (PPF) of fEPSPs, a surrogate measure for presynaptic release probability (Branco and Staras, 2009), in the presence of picrotoxin (Figure 1G). Significantly reduced PPRs of evoked postsynaptic currents (eEPSCs) were further confirmed by patch-clamp recordings (Figure 1H,I). Elevated excitatory neurotransmission in ATG5-cKO mice, thus, is a presynaptic phenotype that does not appear to be caused by impaired synaptic inhibition. In further support of this we observed an increased frequency (Figure 1J) but insignificantly altered amplitude (Figure S1H) of spontaneous excitatory postsynaptic currents (sEPSCs). Next, we asked whether the observed synaptic phenotype is specific for hippocampal CA1 synapses or represents a more general phenotype. To this aim, we investigated a very different synaptic connection, the hippocampal mossy fiber (mf) synapse, which displays a number of specific features, e.g. low basal release probability, pronounced frequency facilitation, and a presynaptic form of long-term potentiation that lacks NMDA receptor involvement (see (Nicoll and Schmitz, 2005) for review). In addition, a use-dependent amplification of presynaptic Ca^2+^ signaling by axonal ryanodine receptors has been postulated (Shimizu et al., 2008). Indeed, we observed decreased post-tetanic potentiation (PTP) and a block of long-term potentiation (mf-LTP) at hippocampal mossy fiber synapses from ATG5 KO mice (Figure S1I,J). These combined data indicate that loss of neuronal autophagy in the absence of ATG5 causes a gain of synaptic neurotransmission at glutamatergic synapses, both in areas CA1 and CA3 of the hippocampus.

**Figure 2.**
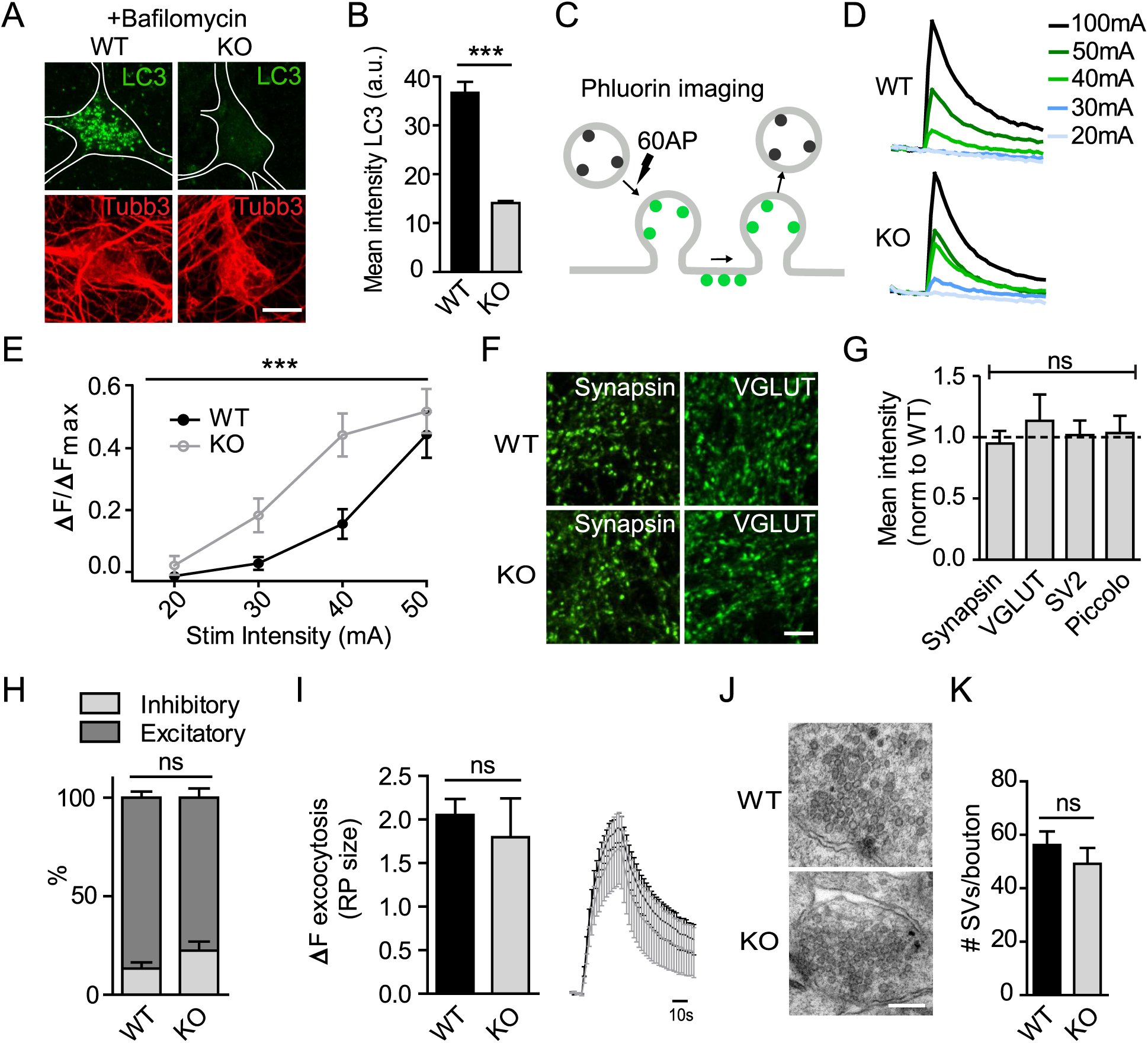
ATG5-iKO neurons display an increased stimulation-dependent SV release. (A,B) Tamoxifen-inducible ATG5-iKO neurons show deficient LC3-positive puncta formation upon bafilomycin treatment (10nm, 4 hours). Representative immunofluorescence images showing LC3 staining in (A) and quantified in (B). Scale bar, 10 μm. n=42 cells, 1 experiment; Mann–Whitney test. (C-E) Detection of exocytosis using Synaptophysin-pHluorin (C) Schematic shows reporter de-acidification during vesicle fusion with the plasma membrane. (D) Example traces showing stimulus dependent decrease in pHluorin signal in a WT and a KO neuron. (E) Graph showing mean peak fluorescence upon different stimulation intensities. Values per cell are normalized to the corresponding maximal fluorescent peak at 100 mA (Fmax). n=17-35 cells, 20 boutons per cell, 4 independent experiments; two-way ANOVA. (F) Representative images for synapsin-1 and VGLUT1 immunostaining of hippocampal neurons. Scale bar, 5 μm (G) Quantification of synapsin-1, VGLUT1, SV2 and Piccolo immunostaining intensities. The mean values for the control are set to 1, and the mean value for the KO is expressed relative to this. n=3 independent experiments, 30 images per condition; one-sample t-test. (H) Percentage of inhibitory and excitatory synapses in WT and KO hippocampal cultures determined by synapsin (marker for all synapses) and vGAT (inhibitory synapse marker) antibody staining. Excitatory synapses are Synapsin-positive and vGAT-negative. n=3 independent experiments, 45-47 images per condition; paired t-test. (I) Quantification and average traces of Synaptophysin–pHluorin expressing neurons stimulated with 600 APs (20Hz) to determine the size of the recycling SV pool (RP). n=3 independent experiments, 20-24 cells per condition; paired t-test. (J) Representative electron micrographs of nerve terminals in WT and KO hippocampal cultures show no difference in number of SVs per bouton. Scale bar, 1 μm. Quantified in (K), n=41 (WT) n=45 (KO) boutons, 1 experiment; Mann–Whitney test. All data represent mean ± SEM, ns: not significant, p < 0.001

**Figure 3.**
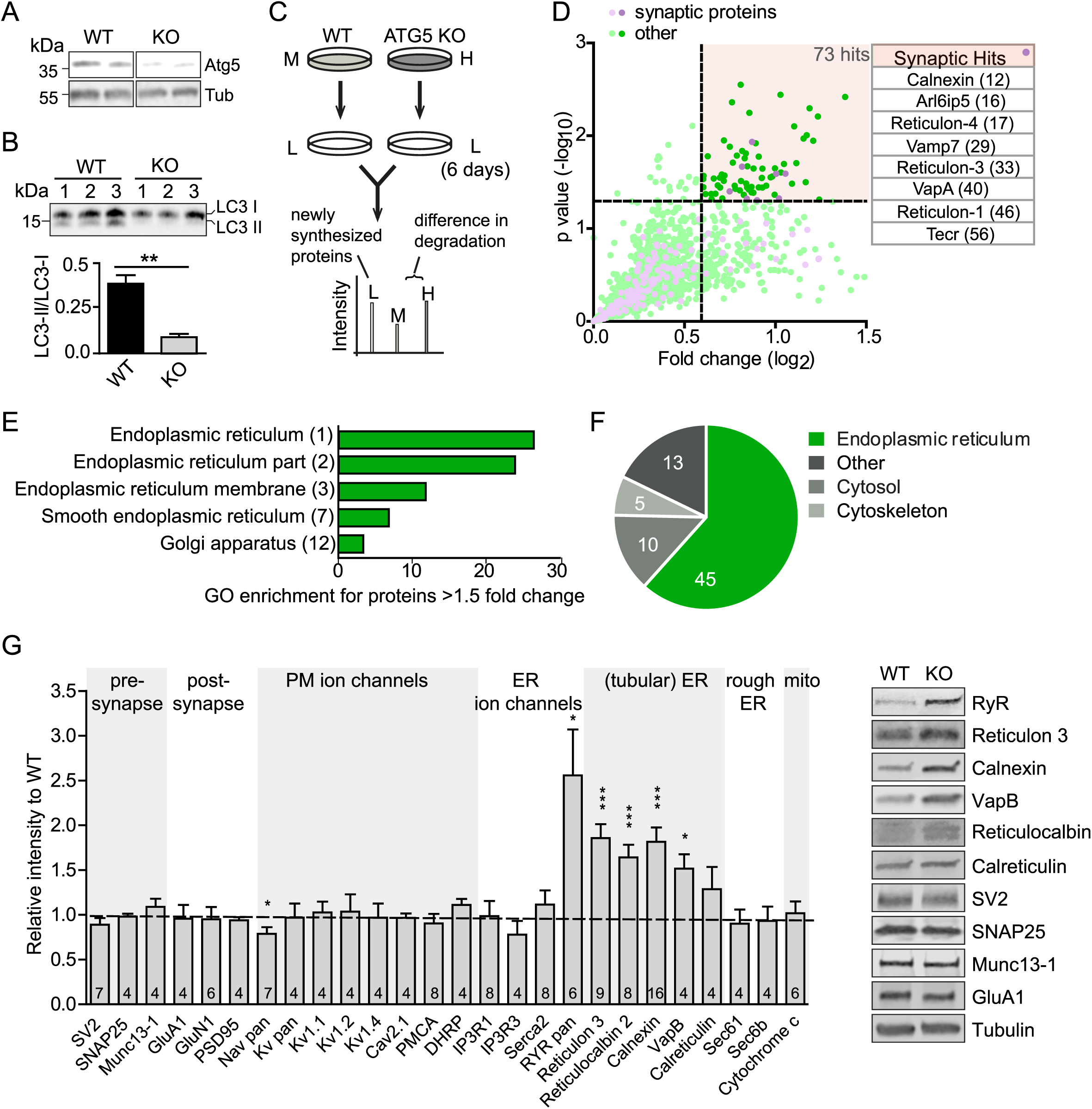
Decreased degradation and accumulation of ER proteins in ATG5-iKO neurons. (A) Immunoblots showing ATG5 decrease in cell lysates from ATG5-iKO neuron cultures compared to control culture lysates. (B) Analysis of LC3-II/ LC3-I ratio in immunoblots of cell lysates from WT and KO neuron cultures. n=3; unpaired t-test. (C) Schematic showing pulsed SILAC procedure to measure protein degradation. Cells were grown for 2 weeks in media containing either heavy or medium variants of lysine and arginine. At day 14, the media was replaced with normal media containing unlabeled (L) amino acids. After 0 (t=0) or 6 (t=6) days cells were harvested, mixed and analyzed by MS analysis, resulting in a list of H/M ratios for each protein. Example is showing an H labeled peptide that is degraded at slower rates than the M labeled peptide, resulting in H/M ratios > 1. (D) Four separate experiments were performed in which 1753 proteins were identified that exhibited H/M (KO/WT) ratios in at least 3 experiments (and 2 conditions, t=0 and t=6). Of the 1753 proteins, 180 are considered to be synaptic proteins. To evaluate protein degradation over the course of 6 days, ratios at t=6 are divided with t=0 ratios and fold changes are plotted. Of 1753 proteins, 73 showed a significant increase in average H/M ratios over the period of 6 days (defined as log_2_fold change>0.6 and p<0.05, dotted lines), that is, exhibited slower degradation rates in the ATG5-iKO neurons. Table shows the protein hits considered to be synaptic (rank of hit is shown in brackets). (E) Gene Ontology analysis indicates that most of the proteins that show slower degradation rates in KO neurons (fold change>1.5) are localized to the endoplasmic reticulum (rank of overrepresented GO Cellular component is shown in brackets). (F) Main subcellular localization of the 73 hit proteins (UniProt-GO Annotation Database) (G) Immunoblot analysis and representative examples of lysates from WT and KO neuron culture, using antibodies against the indicated proteins. Bars show the protein level change of indicated proteins normalized to the housekeeping gene tubulin. The mean values for the controls are set to 1. n= indicated in bars; one-sample t-test. All data represent mean ± SEM, ∗p < 0.05; ∗∗p < 0.01; ∗∗∗p < 0.001.

We challenged these unexpected findings in slices by optical imaging experiments in cultured neurons. To this aim we crossed ATG5*^flox/flox^* mice with a transgenic CAG-*iCre* line in which Cre recombinase activity is under tamoxifen control. We then prepared primary neurons from the hippocampus of these ATG5*^flox/flox^*; CAG-*iCre* (referred to as ATG5-iKO hereafter) mice and corresponding wild-type (WT) mice and treated them with tamoxifen to acutely disrupt the ATG5 gene. As expected, tamoxifen-induced conditional loss of ATG5 in hippocampal neurons abrogated the formation of LC3-containing autophagosomes (Figure 2A,B). To study the effects of defective autophagy in absence of ATG5 on presynaptic function, we monitored SV exo-endocytosis using pH-sensitive pHluorin as a reporter (Kavalali and Jorgensen, 2014) (Figure 2C). Synaptophysin-pHluorin-expressing hippocampal neurons from WT or ATG5-iKO mice were stimulated with 60 action potentials (APs) at different stimulation intensities and SV exo-endocytosis was monitored by optical imaging (Figure 2D). Lower stimulation intensities were required in ATG5-iKO neurons to elicit Synaptophysin-pHluorin responses with maximal amplitudes (Figures 2E, S2A), akin to our electrophysiological data from acute slice preparations (compare Figure 1D). No significant alterations in the levels or localization of SV proteins such as Synapsin 1, SV2, or the vesicular glutamate transporter (vGLUT1), or of the active zone protein Piccolo were detected (Figure 2F,G). Moreover, acute genetic loss of neuronal autophagy in ATG5-iKO neurons did not affect the ratio of excitatory *vs*. inhibitory synapses (Figure 2H), the number of releasable SVs (Figure 2I), or the total SV pool size at the ultrastructural level (Figure 2J,K). Hence, loss of neuronal autophagy causes a cell autonomous facilitation of presynaptic neurotransmission that is not explained by alterations in presynaptic vesicle numbers, pool size, or localization.

### Accumulation of axonal tubular ER induced by block of neuronal autophagy in the absence of ATG5

As enhanced excitatory neurotransmission did not appear to be caused by the accumulation of presynaptic exo- or endocytic proteins or of SVs, we conducted an unbiased quantitative proteomic analysis of the steady-state levels and turnover of neuronal proteins in WT *vs*. ATG5-iKO neurons to identify factors that conceivably might regulate neurotransmission. We treated WT or ATG5-iKO cerebellar granule neurons (CGNs) cultured in the absence of astrocytes with tamoxifen to induce ATG5 loss (Figure 3A), resulting in a block of autophagosome formation as evidenced by defective conversion of the key autophagy component LC3 from its inactive LC3-I to the active LC3-II isoform (Figure 3B). We then conducted quantitative proteomic analysis of neuronal protein turnover by stable isotope labeling with amino acids in cell culture (SILAC) experiments. CGNs were grown in media containing heavy or medium variants of lysine and arginine for 14 days and either analyzed directly by tandem mass spectrometry (MS/MS) to determine their steady-state levels or pulsed for a further 6 days in media containing light (i.e. unlabeled) amino acids before MS/ MS analysis (Figure 3C). Out of the 1,753 proteins identified in at least 3 out of 4 experiments (Table S1), 73 proteins exhibited a reduced degradation rate evidenced by a significantly increased ratio of heavy (KO)- to-medium (WT)-labeled peptides (H/M ratio) over the 6 day-period (i.e. increased (H/M) t=6 / (H/M) t=0) including several allegedly synaptically localized (Hakim et al., 2016) ER membrane proteins (i.e. Reticulon-1, Reticulon-4, VapA, Calnexin) (Figure 3D, Table S1). Many of these factors displayed increased levels already at steady-state (Figure S2B,C). Further gene ontology analysis indicated that the majority of proteins with reduced turnover in the absence of ATG5-mediated neuronal autophagy were proteins known to be localized to the ER (Berner et al., 2018; Saheki and De Camilli, 2017; Westrate et al., 2015) with a preference for tubular ER membrane proteins (Figure 3E,F). To confirm these data by an independent approach, we determined the steady-state levels of distinct classes of ER membrane proteins (i.e. tubular *vs.* rough/ sheet ER) by quantitative immunoblotting. This analysis revealed a prominent accumulation of tubular ER membrane proteins such as Reticulon 3, VapB, and the ryanodine receptor (RyR), an ER-localized ligand-gated calcium channel (Del Prete et al., 2014) (Figures 3G, S2D for reduced degradation rate). Lumenal ER proteins such as Reticulocalbin and Calreticulin were moderately accumulated, while no change in the levels of rough ER membrane proteins such as Sec61and Sec61b involved in secretory protein synthesis was detectable (Figure 3G). Strikingly, we observed no change in the levels of presynaptic vesicle (i.e. SV2) and active zone proteins (i.e. Munc13-1), postsynaptic (i.e. GluA1, GluN1) and plasma membrane ion channels including voltage-gated calcium- (i.e. Cav2.1) or K^+^-channels (i.e. Kv1.1, Kv1.2) and their associated factors, or in mitochondrial cytochrome c (Figure 3G). These data indicate that the tubular ER is a major substrate for neuronal autophagy mediated by ATG5 in healthy unperturbed neurons in the absence of proteotoxic challenge.

Previous work by live imaging has established that autophagosomes preferentially form in distal axons and at presynaptic sites (Hill and Colon-Ramos, 2020; Maday and Holzbaur, 2014; Maday et al., 2012) via a largely constitutive mechanism (Maday and Holzbaur, 2016) that depends on ATG5. We therefore studied whether the accumulation of tubular ER detected at the proteomic level was homogenous throughout the neuron or specific to axons *vs.* the neuronal soma or dendrites. Confocal imaging of hippocampal neurons from tamoxifen-treated ATG5-iKO mice revealed a pronounced accumulation of tubular ER marked by Reticulon 3 in Tau-positive/ MAP2-negative axons, while no change in tubular ER levels was observed in neuronal somata or in dendrites (Figure 4A,B). In ATG5-iKO neurons the axonal ER often appeared as distinct varicosities, possibly representing accumulated ER tubules (Figure 4A and below). Similar results were seen if the ER was marked by Calnexin (Figure S3A) or upon transfection with dsRed-KDEL, a probe for the ER lumen (Figure S3B,C). Loss of ATG5 in astrocytes did not result in accumulated ER in axons (Figure 4D), indicating that the observed neuronal ER phenotype is cell-autonomous. ER-containing axonal varicosities were clearly distinct from p62-containing Ubiquitin conjugates detected mostly in neuronal cell bodies of ATG5-iKO neurons (Figure S3D). To verify that the tubular ER accumulation in axons indeed is a consequence of perturbed neuronal autophagy rather than a phenotype unique to ATG5 loss, we acutely blocked neuronal autophagy by inhibiting VPS34, a phosphatidylinositol 3-phosphate-synthesizing lipid kinase required for the early steps of autophagy (Ariosa and Klionsky, 2016; Ravikumar et al., 2008; Vijayan and Verstreken, 2017). Acute pharmacological inhibition of VPS34 by an established specific small molecule inhibitor, VPS34-IN1 (Bago et al., 2014; Ketel et al., 2016), phenocopied genetic loss of ATG5 with respect to the accumulation of tubular ER in axons (Figure S3,E). Moreover, tubular ER marked by Reticulon 3 also accumulated in Tau-positive axons, often as punctate varicosities, in hippocampal neurons depleted of the early-acting autophagy protein FIP200 by lentiviral knockdown (Figure 4C). In contrast, loss of ATG5 did not affect the levels or localization of the Golgi complex or LAMP1 containing late endosomes/ lysosomes (Figure S3F), the rough ER marked by Sec61b (Figure S3G), or of mitochondria (Figure S3H,I). In spite of the pronounced accumulation of axonal ER, no signs of an induction of the ER stress response probed by specific antibodies against the active phosphorylated form of the ER stress-induced kinases PERK or JNK were detectable in ATG5-iKO neurons (Figures 4E,F, S3J). Moreover, ER tubule diameter, a surrogate measure for ER stress (Schuck et al., 2009; Zhang and Hu, 2016), analyzed by electron microscopy (EM) was unchanged in ATG5-iKO neurons (Figure S3K).

**Figure 4.**
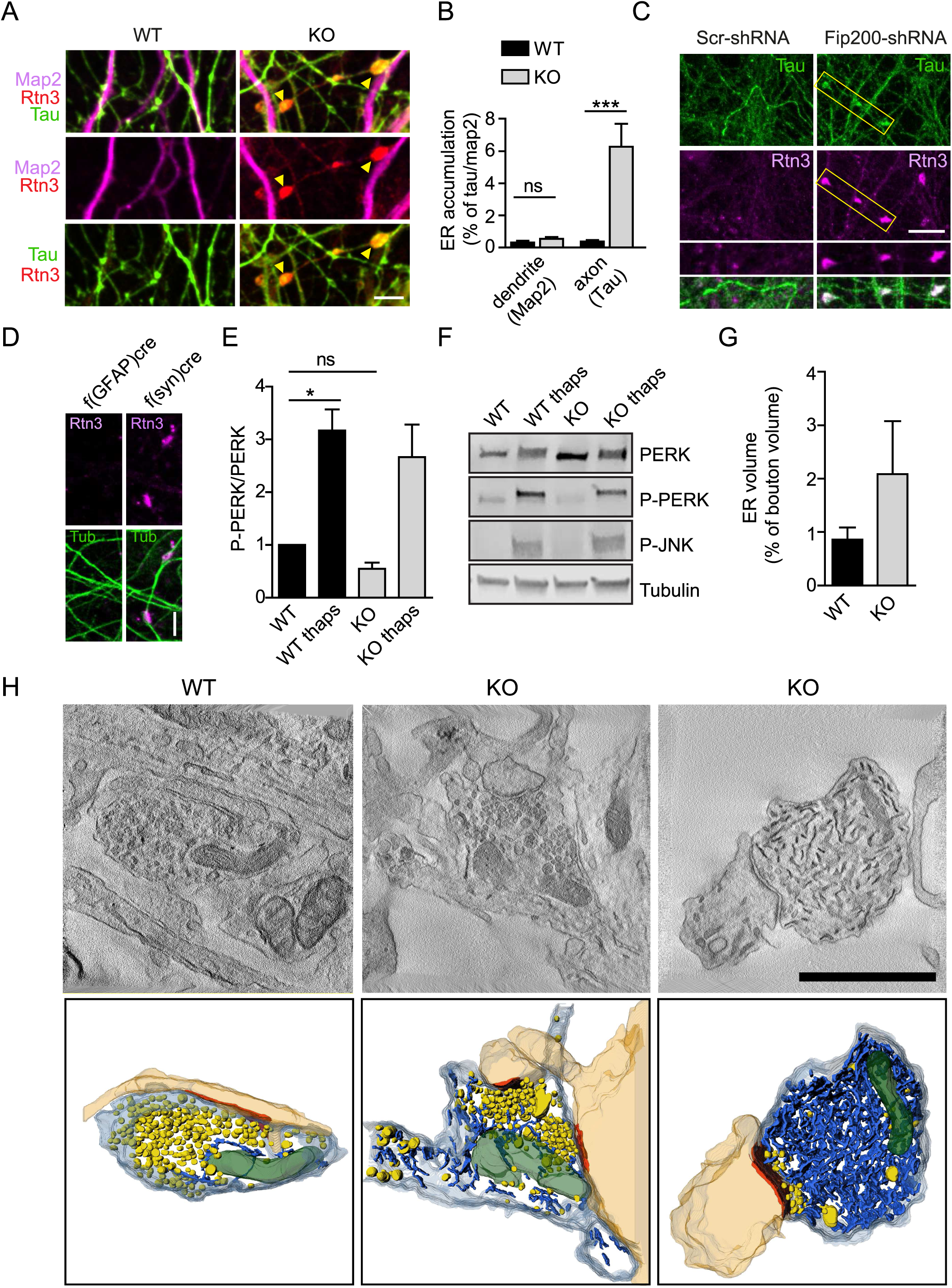
Inhibition of neuronal autophagy leads to accumulation of smooth ER proteins and tubules in axons. (A) Representative confocal images of WT and ATG5-iKO hippocampal neurons immunostained for ER marker Reticulon 3 (Rtn3), axonal marker Tau and dendritic marker Map2. Yellow arrows indicate ER accumulations in KO axons. Scale bar, 5 μm. (B) Quantification of Rtn3 in WT and KO neurons, expressed as dendritic (Map2) or axonal (Tau) area covered with Rtn3 accumulations. n=50 images, 4 independent experiments; Mann– Whitney test. (C) Representative confocal images of control hippocampal neurons and neurons transduced with Fip200-shRNA immunostained for ER marker Rtn3 and axonal marker Tau. ER accumulates in Fip200 KD axons. Scale bar, 10 μm. (D) Cre-mediated ATG5 depletion in neurons (f(syn)cre) but not in astrocytes (f(gfap)cre) leads to neuronal Rtn3 accumulation. Scale bar, 10 μm. (E,F) No basal activation of ER stress pathways in ATG5-iKO neurons. (E) Quantification of PERK phosphorylation (p-PERK) in WT and ATG5-iKO neurons, with and without thapsigargin treatment to induce ER stress (thaps, 1 μM, 16 hours). Conditions are compared to WT, values for WT were set to 1. n=4 experiments; one-sample t-test. (F) Representative immunoblots of lysates from WT and KO neuron cultures, showing PERK, p-PERK, p-JNK and tubulin content. See also Figure S3K for quantifications. (G,H) 3D analysis of of ER tubules in synaptic terminals. (G) Bars indicate the average ER volume in WT and ATG5-iKO boutons. n=12 tomographic reconstructions, 12 boutons per group. (H) Single virtual sections and 3D TEM tomography reconstructions of synaptic boutons showing postsynaptic densities (orange), ER tubules (blue), synaptic vesicles (yellow) and mitochondria (green). Examples are showing a WT bouton and two ATG5-iKO boutons with medium and severe ER volume increases. Scale bar, 1 μm. All data represent mean ± SEM, ns: not significant; p < 0.05; p < 0.001.

The data thus far suggest that block of neuronal autophagy in absence of ATG5 causes the accumulation of tubular ER in axons and, possibly, at synapses. We probed this hypothesis at the ultrastructural level by electron tomography. Tomographic analysis confirmed the dramatic accumulation of ER tubules in axons and at presynaptic sites (Figure 4G,H). Elevated numbers of ER tubules were observed at ATG5-iKO boutons (Figure 4G, 4H/ middle panels). In some cases presynaptic boutons were filled with ER tubules (Figure 4H/ right panels), suggesting that neuronal autophagy is preferentially active in a subset of nerve terminals and/ or distal axons.

Collectively, our findings show that block of neuronal autophagy in the absence of ATG5 causes a pronounced accumulation of tubular ER in axons and at presynaptic sites, whereas the core machinery for neurotransmission and SV exo-endocytosis appears to be unperturbed.

### Accumulation of tubular ER in axons of ATG5 KO neurons is caused by a selective block of autophagy/ lysosome-mediated turnover of ER membranes

We hypothesized that the accumulation of axonal ER under conditions of ATG5 loss is a consequence of defective autophagy/ lysosome-mediated turnover of tubular ER in axons, a process referred to as ER-phagy (Grumati et al., 2018; Khaminets et al., 2015; Liang et al., 2018). We first probed this by inhibiting lysosomal proteolysis by application of the v-ATPase inhibitor bafilomycin in astrocyte-free CGN cultures from WT of ATG5-iKO mice. Bafilomycin treatment of WT neurons for 24h resulted in the accumulation of ER membranes marked by Calnexin. In contrast, bafilomycin failed to cause a further elevation of Calnexin-positive ER membranes in ATG5-iKO neurons (Figure 5A,B), suggesting that ER accumulation in ATG5 KO neurons indeed is the result of defective autophagy/ lysosome-mediated ER degradation. Consistent with these biochemical data, we found the ER tagged with dsRed-KDEL to efficiently co-traffic with LC3-eGFP-containing autophagosomes in distal axons of neurons from WT (Figure 5E) but not from ATG5-iKO mice (Figure 5C,D). No co-transport of dsred-KDEL-labeled ER membranes with LC3-eGFP-containing autophagosomes was observed in dendrites (Figure 5F). Furthermore, recruitment of endogenous LC3 to tubular ER membranes in the axon was observed upon acute pharmacological block and subsequent washout of VPS34-IN1 to reversibly induce neuronal autophagy (Figures 5G, S4A). These data indicate that the axonal ER is a prominent substrate of neuronal autophagy, eventually resulting in ER turnover in the neuronal soma, where most lysosomes reside. We directly tested this hypothesis using a recently developed biosensor for ER membrane turnover via autophagy, i.e. ER-phagy (Liang et al., 2018). This sensor monitors the lysosomal delivery of a chimeric reporter comprised of the pH-sensitive fluorescent protein eGFP (i.e. a probe quenched upon delivery to acidic lysosomes) and pH-insensitive mCherry fused to the ER membrane protein RAMP4. When expressed in WT hippocampal neurons eGFP-mCherry-RAMP4 exhibited a reticular staining pattern, consistent with its ER localization, as well as distinctive mCherry-containing red fluorescent puncta corresponding to ER-containing acidic lysosomes. Such red fluorescent ER-containing acidic lysosomes were rarely observed in ATG5 KO neurons, consistent with a defect in ER-phagy caused by neuronal loss of ATG5. Defective ER-phagy was rescued by re-expression of ATG5 (Figure 5H,I). Surprisingly, loss of ATG5 did not affect autophagic turnover of mitochondria (i.e. mitophagy) (Figures 5J, S4B), consistent with data showing that ATG5 may be dispensable for mitophagy (Honda et al., 2014; Nishida et al., 2009). We conclude that the accumulation of axonal ER under conditions of ATG5 loss is a direct consequence of impaired autophagy/ lysosome-mediated turnover of tubular ER in axons.

**Figure 5.**
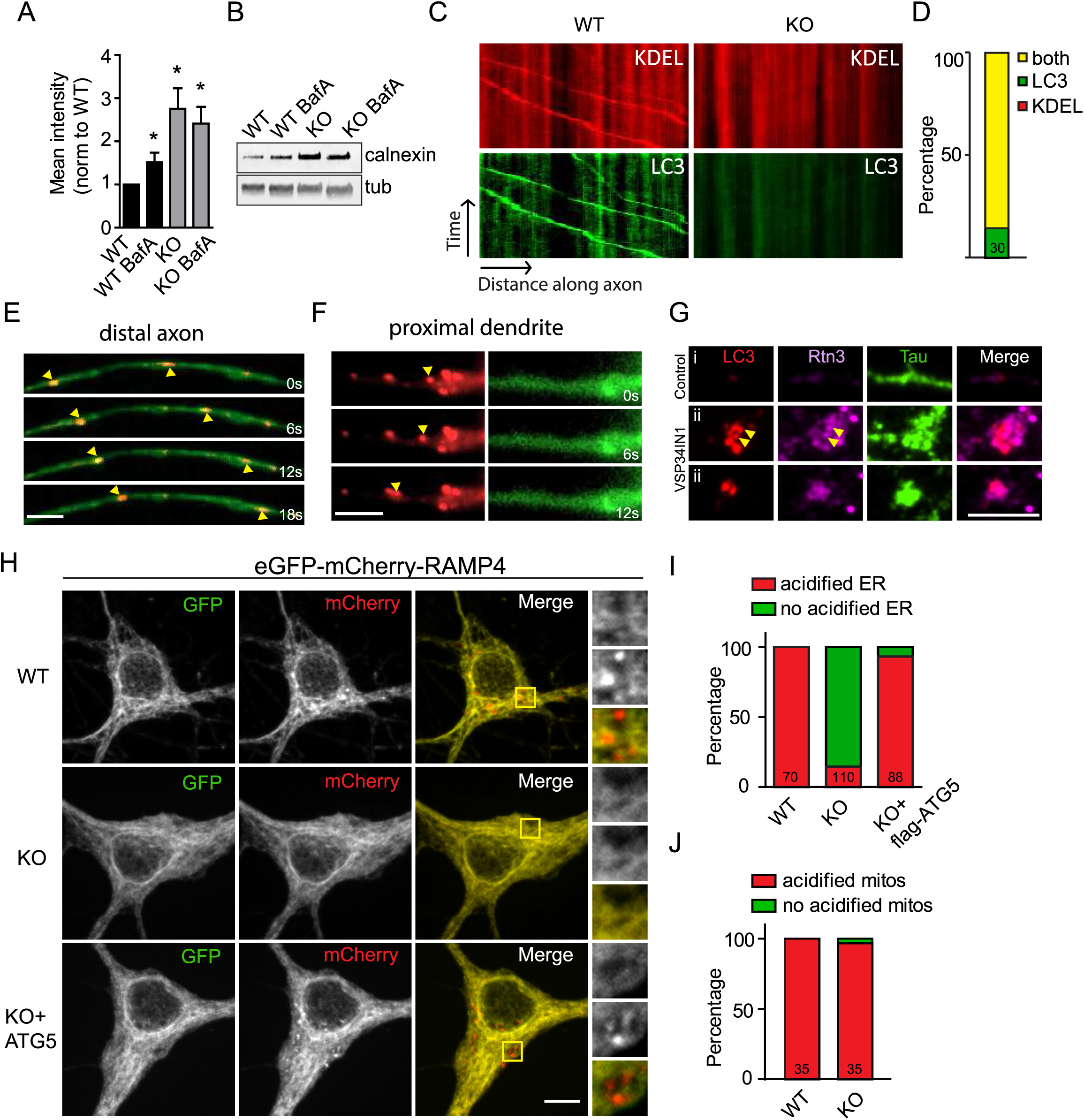
Neuronal ER co-traffics with axonal autophagosomes and is degraded in lysosomes. (A,B) Inhibiting autophagy by preventing vacuolar acidification leads to calnexin accumulation (A) Quantification of immunoblots of WT and ATG5-iKO neurons treated with Bafilomycin A (BafA, 2 nM, 24 hours). All conditions are compared to WT, values for WT were set to 1. n=7 (WT, WT BafA, KO) or 6 (KO BafA) experiments; one-sample t-test. (B) Representative immunoblots of lysates from WT and ATG5-iKO neuron treated with Bafilomycin A. (C-F) Axonal ER co-traffics with LC3-positive autophagosomes. (C) Kymographs showing the colocalization and cotransport of DsRed-KDEL with GFP-LC3b-labelled autophagosomes in WT axons. (D) The majority, but not all, of GFP-LC3b-labelled autophagosomes are positive for DsRed-KDEL. (E, F) Representative time series of colocalization and cotransport of DsRed-KDEL with GFP-LC3b-labelled autophagosomes in an axon (E) Representative time series of a proximal dendrite (F). Yellow arrows indicate moving DsRed-KDEL vesicles. (G) Neurons immunostained for endogenous LC3, Rtn3 and Tau. Blocking autophagy by a selective VPS34 inhibitor (1 μM, 24 hours) and subsequent washout (4 hours) results in axonal LC3 puncta formation that are positive for Rtn3 (i: example of non-treated axon, ii: examples of VPS34 inhibitor treated neurons after washout). (H,J) Acidification of neuronal ER in WT but not in ATG5-iKO neurons. (H) Neurons transfected with eGFP-mCherry-RAMP4. eGFP is quenched as a result of low pH, causing a switch from GFP+/mCherry+ to GFP−/mCherry+ during lysosomal degradation of ER. Yellow boxes indicate magnifications shown on the right. (I) GFP−/mCherry+ (acidified, red) RAMP4 is present in WT neurons but not in ATG5 knockout neurons. Acidified ER is present again in ATG5-iKO neurons after co-expression with flag-ATG5. n=6 experiment for WT and KO and n=3 experiments for KO+ flag-ATG5, total number of cells are indicated in bars. (J) ATG5 depletion does not influence acidification of mitochondria, measured by eGFP-mCherry-TOM20. n=3 experiments, total number of cells are indicated in bars. See also Figure S4B. Scale bars, 5 μm. All data represent mean ± SEM, p < 0.05.

### Elevated calcium release from ER stores via ryanodine receptors accumulated in axons and at presynaptic sites facilitates neurotransmission in absence of ATG5-mediated neuronal autophagy

Major functions of the tubular ER are the (i) transfer of phospholipids such as phosphatidylinositol across contact sites with the plasma membrane, and (ii) the regulation of intracellular calcium signaling and homeostasis (Berner et al., 2018; Bezprozvanny and Kavalali, 2020; Saheki and De Camilli, 2017). We failed to detect significant alterations in the levels of phosphatidylinositol 4-phosphate and phosphatidylinositol 4,5-bisphosphate, the major products of plasma membrane lipid kinases that capitalize on substrate supply of phosphatidylinositol from ER membranes (Saheki and De Camilli, 2017), in ATG5 KO neurons (Figure S4C). Moreover, no change in the dynamics of axonal ER lumenal proteins were observed in fluorescence recovery after photobleaching (FRAP) experiments (Figure S4D) as might be expected, if ER membrane integrity and function were compromised. Hence, we followed the alternative hypothesis that the accumulation of tubular ER in axons might cause alterations in calcium homeostasis, and, thereby, facilitate calcium-triggered presynaptic neurotransmission (Bezprozvanny and Kavalali, 2020; Galante and Marty, 2003). We tested this hypothesis by assaying the relative calcium levels in the axoplasm of WT *vs.* ATG5-iKO neurons using Fluo-8 as a reporter. Axoplasmic calcium levels were elevated about two-fold in ATG5-iKO compared to WT neurons (Figure 6A). In contrast, quantitative measurement of the calcium concentration in the axonal lumen of the ER using ER-GCaMP6-150 (de Juan-Sanz et al., 2017) revealed a reduction from 200 mM in WT neurons to about 100 mM in ATG5-iKO neurons (Figure 6B). These data suggest that accumulation of tubular ER in axons of ATG5 KO neurons leads to elevated calcium efflux from the ER lumen into the axoplasm, which might conceivably disturb presynaptic calcium homeostasis. Indeed, when presynaptic calcium buffering in response to sustained train stimulation (50 Hz, 20 s) was probed by lentivirally encoded Synaptophysin-GCaMP6, we found a significantly reduced ability of ATG5 KO neurons to restore steady-state calcium levels (Figure 6C), suggesting a defect in calcium buffering, likely as a consequence of disturbed calcium homeostasis.

**Figure 6.**
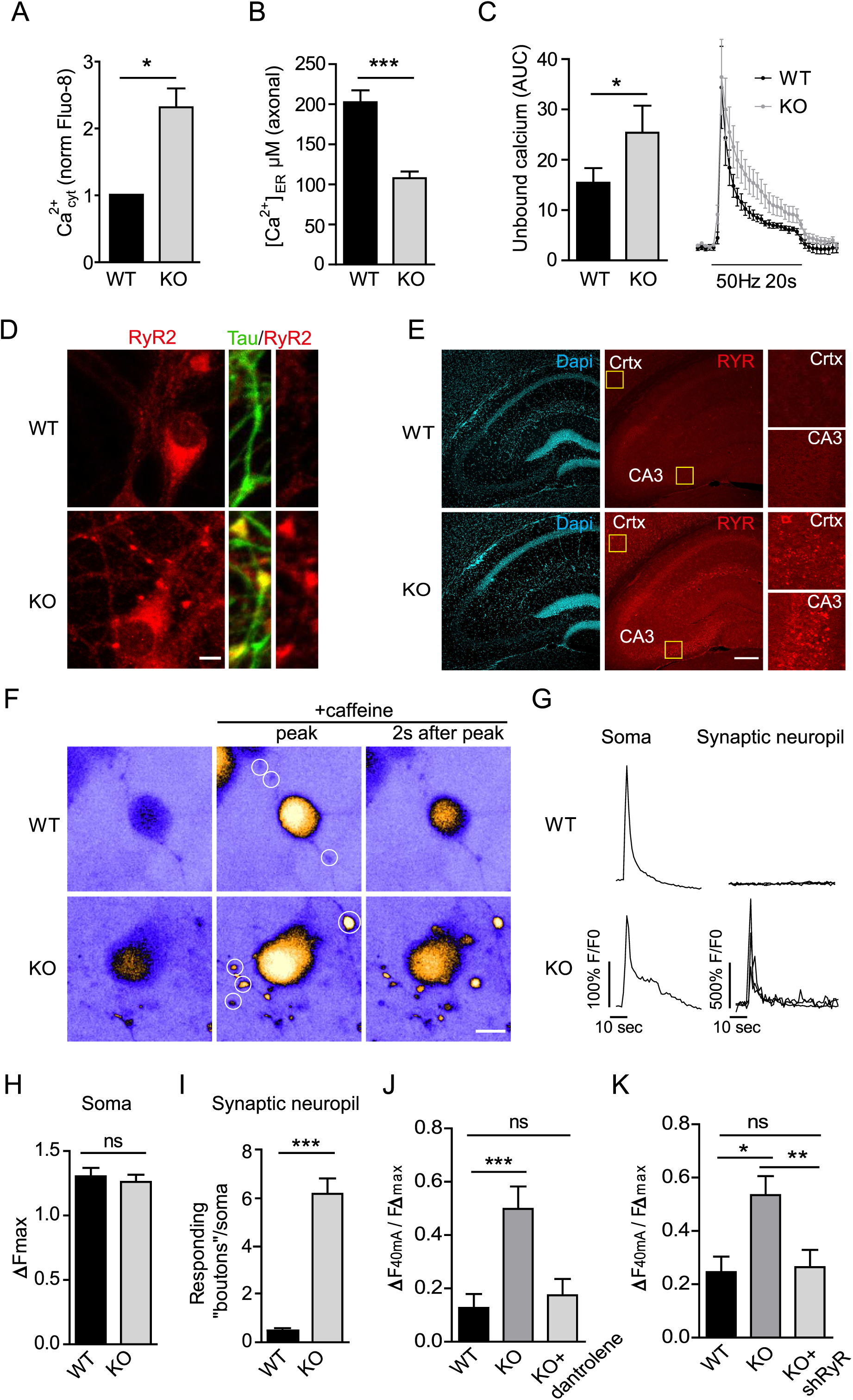
Increased Ryanodine receptor-mediated calcium release underlies elevated neurotransmission in ATG5 KO neurons. (A-C) Impaired calcium homeostasis in ATG5-iKO neurons. (A) Hippocampal neuron cultures were probed with the fluorescent Ca2+ binding dye Fluo-8 to measure cytosolic calcium in neurites. Fluorescent intensities of baseline Fluo-8 were quantified. Neurites were identified by a mild electrical stimulation causing a Fluo-8 increase. WT calcium levels per experiment were set to 1. n=4 experiments, 36 images for WT and 38 images for KO; one-sample t-test. (B) Hippocampal neuron cultures were transfected with ER-GCAMP6-150 and axons were imaged before and after 50 μM ionomycin application to induce indicator saturation for calibration. Average peak fold-change in fluorescence during ionomycin application are used to estimate resting ER calcium concentration in the axon. n=30 axons, 3 independent experiments; unpaired t-test. (C) Calcium buffering in the presynapse was measured by infecting neurons with synaptophysin-GCamp6 virus. The fluorescent change in response to a 50Hz, 20sec pulse was measured. Average traces are indicated on the right and the area under the curve (AUC) is plotted on the left. n=5 experiments, 58-63 cells per condition; paired t-test. (D) Hippocampal neuron culture immunostained for endogenous Ryanodine Receptor RyR2 and axonal marker Tau. Scale bar, 5 μm. (E) Images of mouse brain sections showing an increase in Ryanodine Receptor immunoreactivity in ATG5-cKO cortex (crtx) and hippocampal CA3 area. Yellow boxes indicate magnifications shown on the right. See also Figure S5A for quantifications. Scale bar, 200 μm. (F-I) Increased caffeine-induced calcium release from the ER in ATG5-iKO neuron cultures (F) Heat map images showing Fluo-8 calcium responses during a 20mM caffeine pulse. Scale bar, 10 μm. (G) Representative traces from Fluo8 responses in the soma or boutons (indicated by white circles in (F)). (H) Maximum Fluo-8 intensity increase in WT and ATG5-iKO somas. n=100 WT and 98 KO cells, 2 independent experiments; unpaired t-test. (I) Average number of responding boutons per soma. n=90 WT and 95 KO somas, 2 independent experiments; Mann– Whitney test. (J) Detection of exocytosis using synaptophysin-pHluorin in WT and ATG5-iKO hippocampal neurons. Graph showing mean normalized peak fluorescence upon a 40mA stimulation. Dantrolene (10 μM, a Ryanodine Receptor inhibitor, rescues increased responses in ATG5-iKO neurons. Values per cell are normalized to the corresponding maximal fluorescent peak at 100 mA (Fmax). n=18-22 cells, 3 experiments; one-way ANOVA with Tukey’s post-test. (K) Ryanodine Receptor knockdown decreases exocytosis in ATG5-iKO neurons. Values per cell are normalized to the corresponding maximal fluorescent peak at 100 mA (Fmax). n=13 WT or 25-27 KO cells, 4 experiments; one-way ANOVA with Tukey’s post-test. All data represent mean ± SEM, ns: not significant, ∗p < 0.05; ∗∗p < 0.01; ∗∗∗p < 0.001.

Defects in axonal and/ or ER calcium homeostasis might conceivably result from altered calcium entry via voltage-sensitive calcium channels (Cav), calcium efflux via the plasma membrane calcium ATPase (PMCA), influx into the ER via SERCA, or elevated efflux from the ER lumen into the axoplasm via Inositol 1,4,5-Triphosphate Receptors (IP_3_Rs) or Ryanodine Receptors (RyRs) (Del Prete et al., 2014; Jahn and Fasshauer, 2012; Nanou and Catterall, 2018; Neher and Sakaba, 2008; Scullin and Partridge, 2010). Quantitative proteomic and biochemical analysis by immunoblotting revealed a dramatic accumulation of RyRs (Figures 3G, S5B) in ATG5-iKO neurons and ATG5-cKO brains, while the levels of voltage-gated calcium P/Q-channels (Cav2.1), PMCA, SERCA2, or of various IP_3_R isoforms (IP_3_R1, IP_3_R3) were unaltered (Figure 3G). Elevated steady-state levels of RyR in Tau-positive axons and in the forebrain were further confirmed by confocal imaging of ATG5-iKO neurons (Figure 6D) and brain sections from ATG5-cKO mice (Figures 6E, S5A), respectively. Given the established function of RyRs in ER calcium homeostasis and in the modulation of presynaptic neurotransmission (Galante and Marty, 2003; Irie and Trussell, 2017; Unni et al., 2004), we hypothesized that elevated calcium release from ER stores is mediated via RyRs accumulated in axons and at presynaptic sites to facilitate neurotransmission in the absence of ATG5. Consistent with this hypothesis, ATG5 KO neurons displayed a dramatic increase over WT controls in caffeine-induced calcium release via RyRs (Sato and Kamiya, 2011) in the synaptic neuropil but not in neuronal somata (Figure 6F-I). Hence, the axonal accumulation of RyRs causes RyR gain-of-function, i.e. facilitated release of calcium from axonal ER stores, consistent with our calcium imaging data (compare Figure 6A-C). To finally determine whether increased calcium release from lumenal ER stores via RyR gain-of-function causally underlies elevated presynaptic neurotransmission, we targeted RyRs for acute pharmacological or sustained genetic perturbations. Pharmacological blockade of RyRs by Dantrolene, a well-established RyR antagonist, or lentiviral knockdown of RyRs (Figure S5C) rescued elevated presynaptic neurotransmission in ATG5 KO neurons to amplitudes characteristic of WT neurons (Figure 6J,K).

We conclude that elevated calcium release from ER stores via RyRs accumulated in axons and at presynaptic sites facilitates neurotransmission in absence of ATG5-mediated neuronal autophagy.

## DISCUSSION

Our collective data based on the conditional KO of ATG5 in excitatory neurons and quantitative proteomics as well as live imaging and electrophysiology reveal a crucial function for neuronal autophagy in the control of the tubular ER in axons to regulate excitatory neurotransmission via RyR-mediated calcium release from ER stores. This model is supported by several converging lines of evidence: First, we show that loss of neuronal autophagy in the absence of ATG5 facilitates excitatory neurotransmission in acute hippocampal slices (Figure 1) and in cultured hippocampal neurons (Figure 2) by increasing presynaptic release probability. Second, we identify using SILAC-based quantitative proteomic analyses of nearly 2,000 neuronal proteins (Figure 3) combined with biochemical and optical imaging assays (Figure 5) components of the tubular ER, e.g. Reticulons and the RyR, as the major substrates of neuronal autophagy. Strikingly, tubular ER accumulation was specific to axons and presynaptic sites (Figure 4) and was not observed in either the soma or in neuronal dendrites. The compartment specificity of ER accumulation in axons fits well with the observation that autophagosomes form primarily (although not exclusively) in distal axons and at presynaptic nerve terminals (Hill and Colon-Ramos, 2020; Maday and Holzbaur, 2014; Maday et al., 2012; Vijayan and Verstreken, 2017). Additional factors may contribute to the compartment-specific ER phenotype. For example, the peripheral tubular ER is closely linked to microtubule plus end-directed kinesin motors (Westrate et al., 2015; Zhang and Hu, 2016), likely resulting in the effective retention of the tubular ER in axons that display a uniform plus-end-out microtubule polarity pattern. Third, we demonstrate that elevated calcium release from ER stores via RyRs accumulated in axons and at presynaptic sites of ATG5 KO neurons facilitates excitatory neurotransmission. These observations are consistent with recent data suggesting major roles for the ER (Bezprozvanny and Kavalali, 2020; de Juan-Sanz et al., 2017; Lindhout et al., 2019) and for calcium release via RyRs in the control of presynaptic neurotransmission (Galante and Marty, 2003; Scullin and Partridge, 2010; Shimizu et al., 2008) and presynaptic forms of synaptic plasticity, e.g. long-term depression at hippocampal CA3-CA3 synapses (Unni et al., 2004). Facilitated RyR-mediated calcium release from axonal ER stores and the concomitant elevation of glutamate exocytosis may explain neuronal cell death (Hernandez et al., 2018; Wang and Qin, 2010) and the strongly impaired postnatal viability of ATG5-cKO mice *in vivo*. The role of axonal ER-localized RyRs in the calcium-triggered facilitation of presynaptic neurotransmitter release described here and before (Galante and Marty, 2003) appears to be distinct from the postulated function of STIM1, an ER protein known to couple to ORAI in the plasma membrane to mediate store-operated calcium entry (Saheki and De Camilli, 2017), in the local regulation of release probability via a so far unknown mechanism (de Juan-Sanz et al., 2017).

In addition to their function in the regulation of neurotransmitter release ((Galante and Marty, 2003; Unni et al., 2004) and this work) RyRs have been found to be located in close apposition to large conductance voltage-gated plasma membrane BK-channels to rapidly regulate AP burst firing (Irie and Trussell, 2017). It is therefore possible that the observed accumulation of RyRs in the axonal ER of ATG5 KO neurons in addition to its effects on presynaptic release probability and SV fusion, alters AP shape and, thereby, neuronal excitability. Consistent with this hypothesis, it has been recently found that loss of autophagy increases the excitability of striatal spiny projection neurons (Lieberman et al., 2020). Future experiments will need to test this possibility in detail.

Accumulation of the ER with associated neurodegeneration has been observed in CRISPR KO mice constitutively lacking the autophagy regulatory factor WDR45 (Wan et al., 2020). Our findings are consistent with this and further suggest that ER-phagy is a major autophagic process in neurons in the absence of proteotoxic challenges at steady-state. The physiological and pathophysiological importance of ER-phagy in neurons is further underscored by the fact that loss FAM134B, an adaptor for selective autophagy of the reticular sheet ER, causes sensory neuropathy due to neurodegeneration in mutant mice and in humans (Khaminets et al., 2015). A number of other adaptors for different forms of ER-phagy have been identified since (Grumati et al., 2018). Whether any of these adaptor proteins are required for axonal ER-phagy in hippocampal neurons described here is unclear. Our own preliminary data (Figure S5E) argue against this. It is possible that the known ER-phagy adaptors are functionally redundant or that so far unidentified adaptors mediate autophagy of axonal ER-phagy in central nervous system neurons. Alternatively, axonal ER-phagy may be a constitutive process, intimately linked to the formation of autophagosomes in distal axons and at presynaptic sites (Hill and Colon-Ramos, 2020; Maday and Holzbaur, 2014; Maday et al., 2012; Vijayan and Verstreken, 2017) that serves a homeostatic role in coupling presynaptic function to the constitutive turnover of RyR-containing axonal ER membranes.

In addition to the largely constitutive formation of autophagosomes in axons, autophagy has been shown to be induced by various conditions ranging from overexpression of aggregation-prone proteins (Corrochano et al., 2012) and ROS-induced protein oxidation (Hoffmann et al., 2019), to the depletion of AZ proteins required for presynaptic function (Okerlund et al., 2017). While we do not detect alterations in the steady-state levels or half-lives of major exo-endocytic and AZ proteins in ATG5 KO neurons (compare Figure 2), our data are not incompatible with these earlier studies. For example, it has been shown that co-depletion of the giant AZ proteins Piccolo and Bassoon triggers the activation of E3 ubiquitin ligases and of key ATG proteins resulting in the targeting of SV proteins for degradation via the ubiquitin-proteasome system and autophagy (Waites et al., 2013), resulting in compromised synapse integrity. How different types of physiological (e.g. neuronal activity, aging) and pathophysiological stimuli (e.g. protein aggregate formation in neurodegenerative diseases) regulate distinct types of autophagy in different types of neurons (e.g. glutamatergic *vs*. dopaminergic neurons) in the brain and in the peripheral nervous system remains a fruitful area for future studies.

## ACKNOWLEDGEMENTS

We thank Heike Stephanowitz, Sabine Hahn, Delia Loewe and Silke Zillmann for technical assistance. Supported by the European Union (H2020-MSCA, 65504-SYNPT, to M.K.), the European Research Council (ERC synergy grant to D.S.), the Deutsche Forschungsgemeinschaft (DFG, German Research Foundation) under Germany’s Excellence Strategy (EXC-2049– 390688087) and the Reinhart-Koselleck-Program (HA2685/ 13-1 to V.H.), the Leibniz SAW program (SAW-2014-FMP-2 359, to V.H.), and the Bundesministerium fn□r Bildung und Forschung to D.S. (Smartage 01GQ1420B) and V.H. (Smartage 01GQ1420C).

## AUTHOR CONTRIBUTIONS

M.K. conducted all imaging and biochemical experiments in hippocampal neurons and in slices, D.P. performed electron microscopy and tomography. G.K. and A.S. carried out all electrophysiological experiments. E.K. conducted quantitative SILAC-based mass spectrometry analysis. M.K., D.S. and V.H. designed the study aided by E.K., and analyzed data and wrote the manuscript.

## STAR*METHODS

## CONTACT FOR REAGENT AND RESOURCE SHARING

Further information and requests for resources and reagents should be directed to and will be fulfilled by the Lead Contact Volker Haucke.

## EXPERIMENTAL MODEL AND SUBJECT DETAILS

### Animals

All animal experiments were reviewed and approved by the ethics committee of the “Landesamt für Gesundheit und Soziales” (LAGeSo) Berlin) and were conducted accordingly to the committee’s guidelines.

- Health/immune status: The animals have a normal health and immune status. The animal facility where the mice are kept is regularly checked for standard pathogens. The health reports can be provided upon request.
- Mice used for all experiments were naive. No drug tests were done. Mice were housed under 12/12-h light/dark cycle and up to five animals per cage, with access to food and water *ad libitum*.
- Mouse strains and crossings: ATG5f^lox/flox^ (B6.129S-Atg5tm1Myok) mice (Hara et al., 2006) were crossed with a tamoxifen inducible Cre line (Hayashi and McMahon, 2002) to generate ATG5-iKO (ATG^flox/flox^ × CAG-Cre). To delete ATG5 in excitatory neurons in neocortex and hippocampus, ATG5^flox/flox^ mice were crossed with an Emx1-Cre line (Iwasato et al., 2000) generating ATG5^flox/-^ × EMX1-Cre mice (first generation). By mating ATG5^flox/-^ × EMX1-Cre with ATG5^flox/flox^ mice we obtained conditional ATG5^flox/flox^ × EMX1-Cre (ATG5-cKO) mice.
- Sample size estimation: No estimation of simple size was done as sample sizes were not chosen based on pre-specified effect size. Instead, multiple independent experiments were carried out using several biological replicates specified in the legends to figures.
- Gender of subjects or animals: Mice from both genders were used for experiments.
- How subjects/samples were allocated to experimental groups: Littermates were randomly assigned to experimental groups. Multiple independent experiments were carried out using several biological replicates specified in the figure legends.

## METHOD DETAILS

### Electrophysiology

#### Slice preparation and instrumentation

Electrophysiology was performed in slices prepared from 2-3 months-old ATG5lox/lox × EMX1-Cre and corresponding control mice. Slices were prepared in oxygenated (95% O_2_ / 5% CO_2_) dissection artificial cerebrospinal fluid (ACSF) at low temperature (3-4°C) using vibroslicer (Leica, VT 1200S). After preparation slices were recovered in a resting chamber (Harvard apparatus, BSC-PC) containing ACSF at room temperature (22-24°C) for at least 1.5 hour before recordings. Recordings were performed in a chamber (Warner instruments RC-27L) filled with ACSF with a solution exchange of 3-5 ml per min at room temperature. An upright microscope (Olympus, BX61WI) was used for slice positioning and electrode placement. Glass stimulating (1-1.5 MΩ) and recordings (1.5-2.5 MΩ) electrodes filled with ACSF were prepared from glass capillaries (Hilgenberg) using micropipette puller Sutter P-1000 (Sutter Instruments). The data were recorded at a sampling rate of 10 kHz, low-pass filtered at 3 kHz using EPC9 amplifier and analyzed using Patch Master software (Heka Elektronics).

#### Recordings of CA1 fEPSPs

Mice decapitated after cervical dislocation and brain quickly extracted into dissection ACSF containing: 2.5 mM KCl, 1.25 mM NaH_2_PO_4_, 24 mM NaHCO_3_, 1.5 mM MgS0_4_, 2 mM CaCl_2_, 25 mM glucose, 250 mM sucrose (pH 7.35-7.40). 350 µm thick transversal slices containing clearly visible hippocampus were prepared from both hemispheres and collected in a resting chamber filled with resting/ recording ACSF supplemented with 120 mM NaCl instead of 250 mM sucrose. After recovery slices transferred into recording chamber stimulation and recording electrodes placed in a visually preselected area of *stratum radiatum* and slowly advanced until maximum responses were obtained. Electrical stimuli of 0.2 ms duration were delivered at 0.05 Hz at the stimulation intensity which induced approximately 30-50% of the maximum responses as baseline stimuli. After stabile baseline recordings of at least 10 min an input/output stimulus response curves were made as a measure of basal excitatory synaptic transmission. Slopes of the fEPSP were plotted against fiber volley (FV) amplitudes as a function of increasing stimulation intensity. Stimulation intensity was increased until the maximal fEPSP were obtained, defined as a response with superimposed population spike (PS) component on decaying fEPSP responses. In experiments performed with presence of GABAR antagonist Picrotoxin (50 µM), to prevent spontaneous epileptiform activity, we introduced a cut with a sharp blade between CA3 and CA1 regions. Short-term synaptic facilitation was tested by delivering two pulses at time intervals from 10 to 500 ms at a stimulation intensity which induced one third of the maximal responses. Paired pulse facilitation (PPF) was calculated as a percentage increase of the amplitude of the second response as compared to the first. For short intervals (10 and 20 ms), the first fEPSPs were digitally subtracted before measurements of the second fEPSPs. Each trace measured for the stimulus response curve and paired pulse parameters is an average of 3 consecutive stimulations delivered every 20 and 30 s for stimulus response curves and paired pulse protocols, respectively.

#### Recordings of MF-fEPSPs

Mice anesthetized with isoflurane and transcardially perfused with ice cold dissection ACSF containing the following substances: 75 mM sucrose, 25 mM glucose, 87 mM NaCl, 25 mM NaHCO_3_, 2.5 mM KCl, 1.25 mM NaH_2_PO_4_, 0.5 mM CaCl_2_, 7 mM MgCl_2_, pH 7.35-7.4. Dissection ACSF was cooled down in a freezer and bubbled at least 30 min prior to use with 95% O_2_ / 5% CO_2_. After 2 minutes of perfusion brain quickly removed and fresh 350 μm-thick hippocampal sections were prepared from both hemispheres and kept in sucrose based cutting/storage solution for recovery at 35 °C for 30 minutes as described in (Bischofberger et al., 2006). Slices were transferred in a resting chamber filled with recording ACSF of following composition: 120 mM NaCl, 2.5 mM KCl, 1.25 mM NaH_2_PO_4_, 25 mM NaHCO_3_, 1.5 mM MgS0_4_, 2.5 mM CaCl_2_, 25 mM glucose, pH7.35-7.4, at room temperature for at least an hour before the use. Mossy fibers (MF) were stimulated in the area of internal side of granule cell layer of the dentate gyrus and MF-*fEPSPs* were recorded in the str. lucidum of the CA3 field. MF-CA3 responses are characterized with the strong presynaptic facilitation and were identified using frequency facilitation parameter in which stimulation frequency is set to 0.3 Hz. The responses which exhibit at least 200% facilitation were accepted as MF-fEPSPs and were recorded further. Basal stimulation were applied every 30 sec in order to monitor stability of the responses at least for 15 minutes before LTP recordings. The stimulation intensity for FF and LTP experiments were selected to 50-60 % and 5 HFS delivered every 30 seconds each one containing 100 pulses at 100Hz were applied to induce LTP. LTP at this synapse can be generated presynaptically and is known to be NMDA receptor-independent, therefore 50 µM APV was bath applied during recordings. In order to confirm that fEPSPs were generated by the stimulation of MFs an agonist of type II metabotropic glutamate receptors DCG IV (2 µM) was applied and only responses inhibited by 70-80% and more were assumed to be elicited by mossy fiber synapses.

#### Whole cell recordings

Slices were recorded in a submerged recording chamber and were perfused with ACSF at a flow rate of 5ml/min. Whole-cell recordings were performed with a K-gluconate–based intracellular solution containing (in mM) K-gluconate (120), HEPES 20, KCl 3, NaCl 7, MgATP 4, NaGTP (0.3), and phosphocreatine 14, adjusted to pH 7.3 with KOH. Paired pulse ratio (PPR) was detected by Schaffer collateral stimulation with a low resistance glass electrode in str. radiatum of CA1. Paired stimulation (50 ms ISI) was applied and the amplitude of the second EPSC was devided by the first EPSC amplitude. Cumulative distribution of PPR was analyzed using 10 PPRs per cell.

Spontaneous EPSCs (sEPSCs) were recorded in voltage clamp configuration and cells were clamped to −60mV. Signals were detected automatically using IGOR Pro with the plugin Neuromatics and subsequently manually sorted by visual inspection. Cumulative distribution of sEPSC interevent interval (IEI) was analyzed using an equal number of events per cell per condition to prevent overrepresentation of single neurons. Only cells where at least 30 IEIs could be detected were taken into account for the distribution.

### Expression constructs, shRNA and lentivirus production

Synaptophysin 1 fused to pHluorin was kindly provided by L. Lagnado (MRC Laboratory of Molecular Biology, Cambridge, UK). ER-GCAMP6-150, TetOn-eGFP-mCherry-RAMP4, TOM20MTS-mCherry-EGFP-Tet-On, sRed2-Mito-7 and pEGFP-LC3 were obtained from Addgene. DsRed-KDEL was created by inserting an ER retention signal sequence (AAGGACGAGCTG) in a pDsRed2 expression vector just before the stopcodon. For viral-mediated expression, lentiviral vectors expressing synaptophysin fused C terminally with GCamp6f controlled by the human synapsin-1 promotor, were used. For viral-mediated knockdown, lentiviral vectors expressing nuclear localized RFP controlled by the human synapsin-1 promotor, and the appropriate shRNA controlled by the U6 promoter, were used. For target and non-target control shRNA sequences see the Key Resources Table. Lentiviral particles were produced by the viral core facility of the Charité – Universitätsmedizin Berlin, Germany. See Key Resources Table for further information.

### Antibodies

See Key Resources Table.

### Neuron preparation, culture, infection, and transfection

Neuronal cultures were prepared by surgically removing the hippocampi or cerebellum from postnatal mice at p1-3 (hippocampus) or p4-7 (cerebellum), followed by trypsin digestion to dissociate individual neurons. 100,000 hippocampal cells were plated as 40 μl drops per poly-L-lysine coated coverslip and 2 mL of plating medium (basic medium (MEM; 0.5% glucose; 0.02% NaHCO3; 0.01% transferrin) containing 10% FBS, 2 mM L-glutamine, insulin and penicillin/streptomycin) was added 1 h after plating. For cerebellar granule cell (CGN) cultures 1.5×106 cells were added directly to poly-L-lysine coated dished containing 2 mL of plating medium. After one day in vitro (DIV1) 1 mL of plating medium was replaced by 1 mL of growth medium (basic medium containing 5% FBS; 0.5 mM L-glutamine; 2% B27 supplement; penicillin/ streptomycin) and on DIV2 1 mL of growth medium was added. AraC was added to the culture medium to limit glial proliferation. For cerebellar granule cell (CGN) cultures 25mM KCl was added to the plating and growth medium. CGN cultures used for the multiplexed SILAC are grown in Neurobasal medium (described in more detail under the Multiplexed SILAC subheading). To initiate homologous recombination in neurons from floxed animals expressing a tamoxifen-inducible Cre recombinase cultured neurons were treated with 0.3 μM (Z)-4-hydroxytamoxifen (Sigma) immediately after plating. When other drugs are added to the growth medium, concentration and duration of treatment are mentioned in the figure legends.

For lentiviral transduction about 5×10^5^ infectious virus units per 35 mm-diameter well were pipetted onto hippocampal neurons at DIV 1 or 2. A non-targeting shRNA control was included in RYR knockdown experiments. For calcium phosphate transfection 6 μg plasmid DNA, 250 mM CaCl2 and water (for each well of a 6-well plate) were mixed with equal volume of 2x HEPES buffered saline (100 μl) and incubated for 20 min allowing for precipitate formation, while neurons were starved in NBA medium for the same time at 37°C, 5% CO2. Precipitates were added to neurons and incubated at 37°C, 5% CO2 for 30 min. Finally, neurons were washed three times with HBSS medium and transferred back into their conditioned medium. For TetOn-eGFP-mCherry-RAMP4/TOM20 expression, 4 µg/ml doxycycline was added at the day of transfection. Live imaging and fixation of hippocampal cultures was conducted at DIV 13–16, CGN cultures were lysed at DIV13-20.

### Immunostaining of hippocampal neurons in culture

Neurons were fixed on DIV 13–16 with 4% paraformaldehyde (PFA)/4% sucrose in phosphate-buffered saline (PBS) for 15 min at room temperature (RT), washed and incubated with primary antibodies in PBS containing 10% normal goat serum (NGS) and 0.3% Triton X-100 (Tx) overnight at 4 degrees. Coverslips were washed three times with PBS (10-min each) and incubated with corresponding secondary antibodies for 1 hour. Finally, coverslips were washed three times in PBS and mounted in Immumount. Alternatively, for LC3 immunostaining, cells were fixed with PFA and permeabilized with digitonin (200 µg/ml) for 15 min before incubating with primary and secondary antibodies in PBS. For lipid stainings, cells were fixed with 2% PFA/2% sucrose/ 1% glutaraldehyde in PBS for 20 min at RT. Neurons were then permeabilised with 0.5% Saponin /1% BSA in PBS for 30 min at RT and incubated with indicated antibodies diluted in 1%BSA/10%NGS in PBS. Neurons were imaged at a resolution of 1,024 × 1,024 on a Zeiss laser scanning confocal microscope LSM710 or a spinning disc confocal microscope (CSU-X1, Nikon) with a 63× oil objective. All acquisition settings were set equally for all groups within each immunostaining. Image processing and quantitative analysis was performed in ImageJ. For quantitative analysis of fluorescent intensities in the soma the total area of the soma was manually selected and measured using ImageJ selection tools. Average intensities of fluorescent puncta (synapses) were measured by centering 9 × 9 pixel (∼1 × 1 μm) regions on maxima determined by ImageJ processing function. For quantifying ER antibody stainings in neurites MAP2 and Tau signal were used as template for a mask, restricting the quantified area to the shape of the dendrites or axons. For quantifiying lipid levels in axons, Synaptobrevin staining signals were used as a mask. Fluorescent areas were determined by applying thresholding and analysed using the ‘Analyze particles’ ImageJ module to determine the number or area of fluorescent spots.

### Immunohistochemistry on brain sections

2-5 months-old ATG5 KO pups and their WT littermates were euthanized by an overdose (i.p.) of Ketamin (120 mg/kg body weight)/Rompun (16 mg/kg body weight) and transcardially perfused with 4% formaldehyde in 0.1 M PBS. Brains were isolated and postfixed in the same solution overnight at 4 °C. After cryoprotection in 20% sucrose, frozen sections (30 μm) were collected in 0.1□M PBS. For immunostaining, corresponding hippocampal sections from WT and KO littermates were processed simultaneously. Sections were blocked for 2 h in 5% normal goat serum and 0.125M PBS with 0.3% Tween (PBST). Tissue was then washed with PBST and incubated in normal goat serum–PBST mixture for 48 h with primary antibodies. After washing, sections were incubated for 16 h with Alexa-conjugated secondary antibodies and Dapi in PBST. Finally, sections were washed, mounted and coverslipped on gelatin-coated glass slides. Sections were imaged at a resolution of 1,024 × 1,024 using a Zeiss laser scanning confocal microscope LSM710 with a 20x objective (dry). All acquisition settings were set equally for sections of all groups within each immunostaining. Image processing and quantitative analysis of fluorescence intensity was performed in ImageJ. Images were quantified by measuring the mean intensity in defined region of interests (ROI). To quantify RYR area images were thresholded and particles analyzed with the analyze function within defined ROIs. Only particles with sizes larger than 4 pixels were selected for analysis.

### pHluorin imaging

To track synaptic vesicle exo-/ endocytosis, neurons transfected with synaptophysin-pHluorin were subjected to electrical field stimulation using an RC-47FSLP stimulation chamber (Warner Instruments) and imaged at 37°C in imaging buffer (170 mM NaCl, 3.5 mM KCl, 0.4 mM KH2PO4, 20 mM N-Tris[hydroxyl-methyl]-methyl-2-aminoethane-sulphonic acid (TES), 5 mM NaHCO3, 5 mM glucose, 1.2 mM Na2SO4, 1.2 mM MgCl2, 1.3 mM CaCl_2_, 10 μM CNQX and 50 μM AP-5, pH 7.4) by epifluorescence microscopes (Zeiss Axiovert 200M or Nikon Eclipse Ti) equipped with a 40Χ oil objective. Images were acquired at 0.5 or 1 Hz frame rate. Quantitative analysis of responding boutons (20 per stimulation) was performed using ImageJ. Fluorescence intensities of responding boutons were corrected for background and photobleaching if nessesary. For the experiments in which stimulation intensities were varied, each cell was subjected to the different stimulation strengths mentioned (e.g. 20mA-30mA-40mA-50mA-100mA). ΔF was obtained by calculating ΔF = [F (data point fluorescence) - F0 (resting fluorescence)]. ΔFmax is ΔF during a 100mA stimulation.

### Photobleaching experiments

For quantitative fluorescence recovery after photobleaching (FRAP) experiments, neurons were transfected as described before, and imaged on a Zeiss laser scanning confocal microscope LSM710 with *ZEN* 2010 *software*. The acquisition was performed with a 63X oil objective, 1024 x 1024 pixels per image and a zoom factor 4.5. After acquiring 10 pre-FRAP images (every 5 seconds), an 80 pixel long ROI on the proximal axon was photobleached with maximal laser power (10 iterations) and a futher 30 images were acquired. To analyze the recovery of fluorescence, the bleached area was selected and background subtracted by subtracting the intensity of an empty, non-bleached area. Recovery R was calculated as R = (I(t)- I(directly after bleaching))/(I(before bleaching)-I(directly after bleaching)), with I denoting total intensity.

### Ca^2+^ imaging

#### Cytosolic Ca^2+^

Hippocampal neuron cultures from WT and ATG5 KO mice were loaded with 2 µM Fluo-8/AM together with 0.02% pluronic for 15 min at 37°C. Prior to imaging, neurons were washed 3 times in imaging buffer (see heading pHluorin Imaging for recipe). For data shown in Figure 6A neurites were identified by a mild electrical stimulation causing a Fluo8 increase. For the caffeine-induced calcium responses (Figure 6F), calcium was omitted from the imaging buffer and images were acquired at 1 Hz frame rate. After correction for background fluorescence, fluorescence intensity was analyzed. Number of responding boutons per soma (Figure 6I) was determined by counting the responding boutons in a 100×100 μm ROI containing a soma.

#### Synaptic Ca^2+^

Neurons were transduced with Synaptophysin-GCamp6 as described before and subjected to electrical field stimulation using an RC-47FSLP stimulation chamber (Warner Instruments) and imaged in imaging buffer. Images were acquired at 1 Hz frame rate.

#### ER luminal Ca^2+^ measurements

Neurons were transfected with ER-GCAMP6-150 as described before, and axons were imaged in imaging buffer before and after (Fmax) addition of 50 μM ionomycin. Knowing the in vitro characteristics of the indicator used (de Juan-Sanz et al., 2017), baseline [Ca2+]ER is calculated using the following equation: [Ca^2+^]ER=Kd((Fr/Fmax-1/Rf)/(1-Fr/Fmax)^1/n^. Kd is the affinity constant of the indicator (150 μM), Fr is the measured fluorescence at rest, Rf is the dynamic range (45) and n is the Hill coefficient (1.6). Fmax values were not corrected for pH changes.

All Ca^2+^ imaging experiments were performed in imaging buffer at 37°C with an epifluorescence microscope (Nikon Eclipse Ti) equipped with a 40Χ oil objective. Quantitative analysis and image processing were performed using ImageJ.

### Electron microscopy and tomography

DIV14 neurons were fixed with 2% glutaraldehyde in PBS. Coverslips were then postfixed with 1% OsO4 and 1.5% potassium hexacyanoferrat (III), stained en bloc with 1% uranyl acetate, followed by dehydration in a methanol gradient, propylene oxide and Epoxy resin infiltration. After polymerization, coverslips were removed and 50 nm sections were cut and contrasted with uranyl acetate and lead citrate for transmission electron microscopy (TEM) and morphometric analysis (SVs). For TEM tomography, 250 nm sections were cut and collected on coated slotted grids with 10 nm gold fiducials. Series of images from +60° to −60° were taken with a 1° step at Tecnai G20 microscope. Etomo/IMOD and Microscopy imaging browser MIB were used to work with 3D volumes and render 3D models of subsynaptic structures.

### Immunoblot analysis of mouse brain extracts and neuron cultures

Brain tissue was homogenized in lysis buffer (20 mM Hepes-KOH, pH 7.4, 100 mM KCl, 2 mM MgCl2, 1% Triton X-100, supplemented with 1 mM PMSF and mammalian protease and phosphatase inhibitor mixture) using a glass teflon homogenizer. Neuron cultures were lysed in RIPA buffer (150 mM NaCl, 1.0% NP-40, 0.5% sodium deoxycholate, 0.1% SDS, 50 mM Tris, pH 8.0) with protease and phosphatase inhibitors. Lysates were incubated 30 min on ice before centrifugation at 17,000g for 10 min at 4 °C and protein concentrations determined by Bradford or BCA assay. Equal concentration of lysates in Laemmli sample buffer were boiled for 5 min. Between 20 and 60 μg protein was resolved by SDS–PAGE and immunoblotting was done on nitrocellulose membranes. Membranes were incubated with the primary antibodies at 4°C overnight. On the next day, bound primary antibodies were detected by incubation with IRDye 680/800CW-conjugated secondary antibodies via the Odyssey Fc Imaging system (LI-COR Biosciences).

### Multiplexed SILAC and mass spectrometry analysis

CGN WT and KO cultures (1.5-1.7×106 cells per culture) were grown in custom-made lysine and arginine free NB (Life technologies) to which “medium” (M) variants D4-lysine/13C6-arginine (Lys4/Arg6) or “heavy” (H) variants 13C615N2-lysine/13C615N4-arginine (Lys8/Arg10) were added. Growth medium consisted of (Lys/Arg) NB medium supplemented with 2% B-27, 0.5 mM L-glutamine, 25mM KCL and penicillin/streptomycin. After 2 weeks, the cultures were gently washed and growth medium was replaced by conditioned medium from “sister cultures” grown in paralel in “light” (unlabeled Lys/Arg) growth medium. Neurons were harvested and lysed after 0, 2 and 6 days and mixed together as pairs of time matched WT and KO sets. To exclude the possibility of a specific labeling type affecting the experimental outcome, the labeling (heavy or medium type) was varied between the WT and KO samples in the four biological replicates. For clarifying purposes in text, figures and legends, the KO is always heavy labeled (H) and the WT is medium labeled (M).

Forty micrograms of protein in Laemmli sample buffer from each time point was separated on 4–15% SDS–PAGE, each lane was then cut into 15 slices, and in-gel tryptic digestion was performed. Tryptic peptides were analysed by a reversed-phase capillary liquid chromatography system (Ultimate 3000 nanoLC system; Thermo Scientific) connected to an Orbitrap Elite mass spectrometer (Thermo Scientific). Identification and quantification of proteins were performed using MaxQuant (version 1.5.1.0) software. Data were searched against the Uniprot mouse protein database. The initial maximum mass deviation of the precursor ions was set at 20 ppm, and the maximum mass deviation of the fragment ions was set at 0.35 Da. Methionine oxidation and the acrylamide modification of cysteine were used as variable modifications. False discovery rates were <1% based on matches to reversed sequences in the concatenated target-decoy database. Proteins were considered if at least two sequenced peptides were identified.

### Data analysis of SILAC

SILAC quantitation is done using the signals of the medium (Lys4/Arg6) and heavy (Lys8/Arg10) labeled peptides, the unlabeled peptides are ignored. Four independent experiments were performed to compare protein degradation in WT vs KO cultures after 6 days of “light” medium. Only proteins with a H/M ratio in both timepoints (t0 and t6) in 3 out of 4 experiments were considered. The plotted fold changes were calculated by dividing H/M(t6) by H/M(t0). Analyses were performed using Microsoft Excel. Synaptic proteins were manually selected using a list of 314 proteins that are either synapse-specific, highly enriched or implicated in synaptic function (Hakim et al., 2016). GO cellular component enrichments were calculated using GOrilla, using a ranked list of proteins with >1.5 fold change (>0.6 log_2_fold) in KO/WT ratio and the total list of 1753 proteins as a reference. The GO subcellular localization of the 73 hit proteins (defined as >0.6 log_2_fold change and p<0.05) was done manually for each hit using the UniProt-GO Annotation Database.

### Experimental Design

A strategy for randomization, stratification or blind selection of samples has not been carried out. Sample sizes were not chosen based on pre-specified effect size. Instead, multiple independent experiments were carried out using several sample replicates as detailed in the figure legends.

## QUANTIFICATION AND STATISTICAL ANALYSIS

### Imaging and biochemistry

Values are always depicted as mean ± SEM. Significance is denoted using asterisks *P<0.05, ** P<0.01, *** < 0.001 and p>0.05 is not significant (ns). Statistical data evaluation was performed using Graph Pad Prism 5 software. One-sample t-tests were used for comparisons with control group values that had been set to 1 for normalization purposes. For comparisons between two experimental groups statistical significance was analyzed by two-sample, two-tailed unpaired or paired Student’s *t*-tests or Mann–Whitney test (as indicated in the figure legends). Pearson’s chi-square test was used to examine Mendelian ratios. Kolmogorov–Smirnov test was performed to compare the distributions of individual genotypes for data shown as cumulative distribution.. For comparisons between more than two experimental groups statistical significance data was analyzed by one-way ANOVA with post-hoc test (as indicated in the figure legends). The number of animals, cell cultures or cells used (n) is stated in the figure legends. SigmaPlot was used for electrophysiological data analyses, presentation and statistical calculations. Data curves were statistically evaluated using ANOVA with repeated measures (significance depicted over a line encompassing the curve) and comparisons of two groups statistical significance was tested using a two-tailed unpaired Student’s t test.

## KEY RESOURCES TABLE

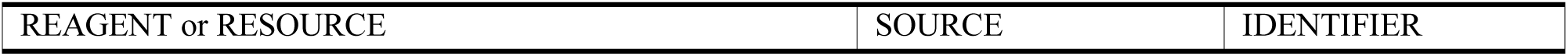
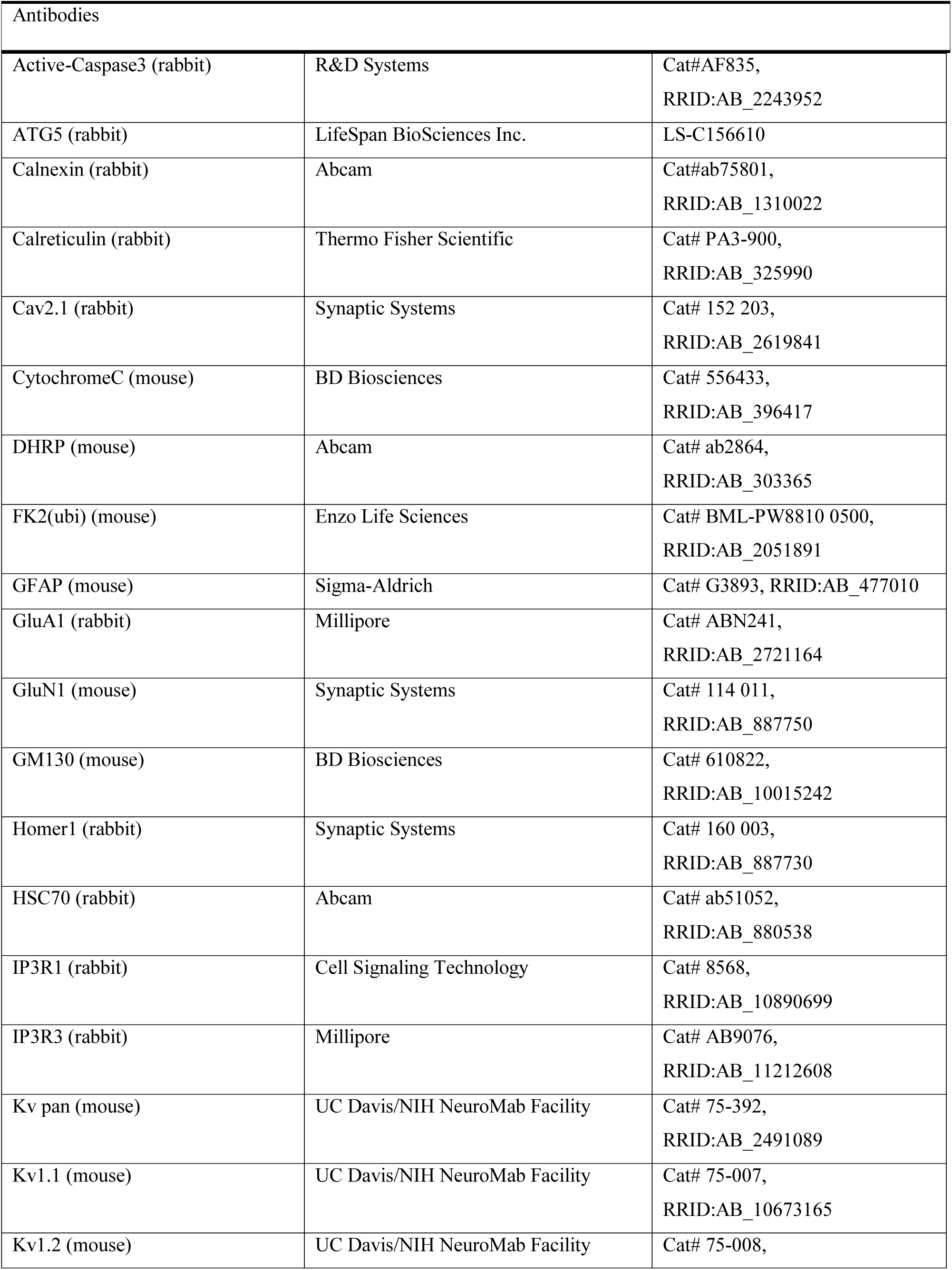
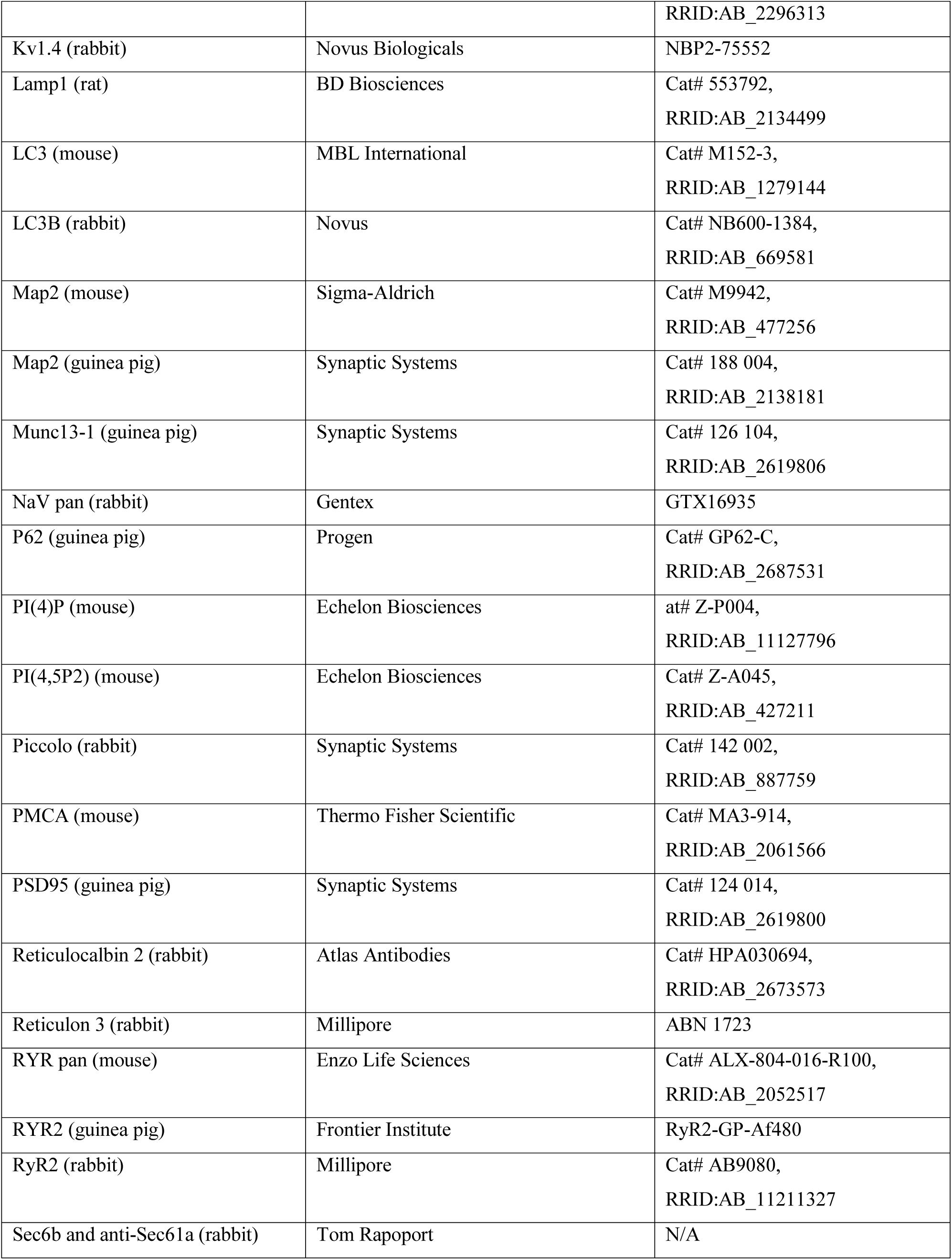
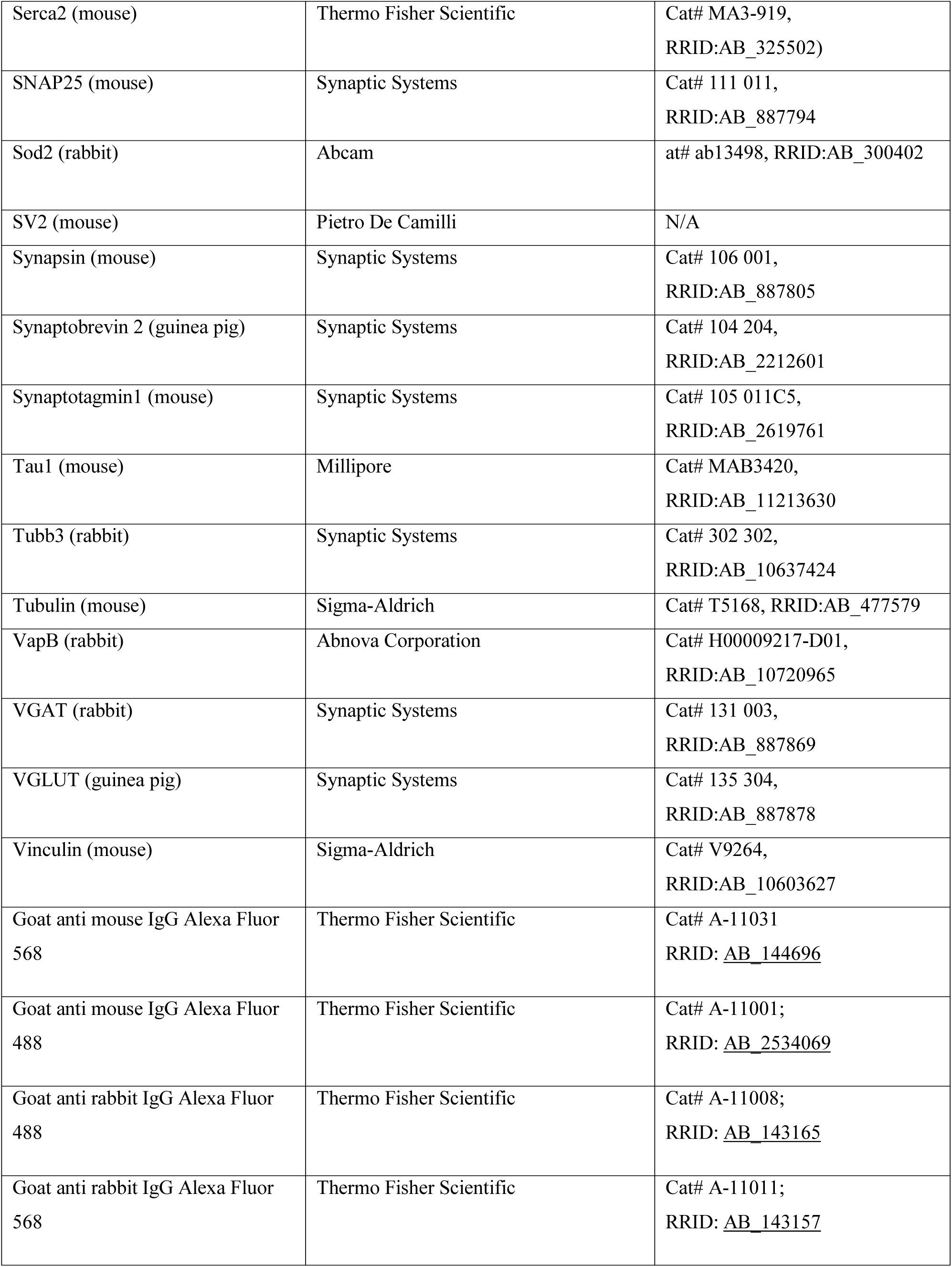
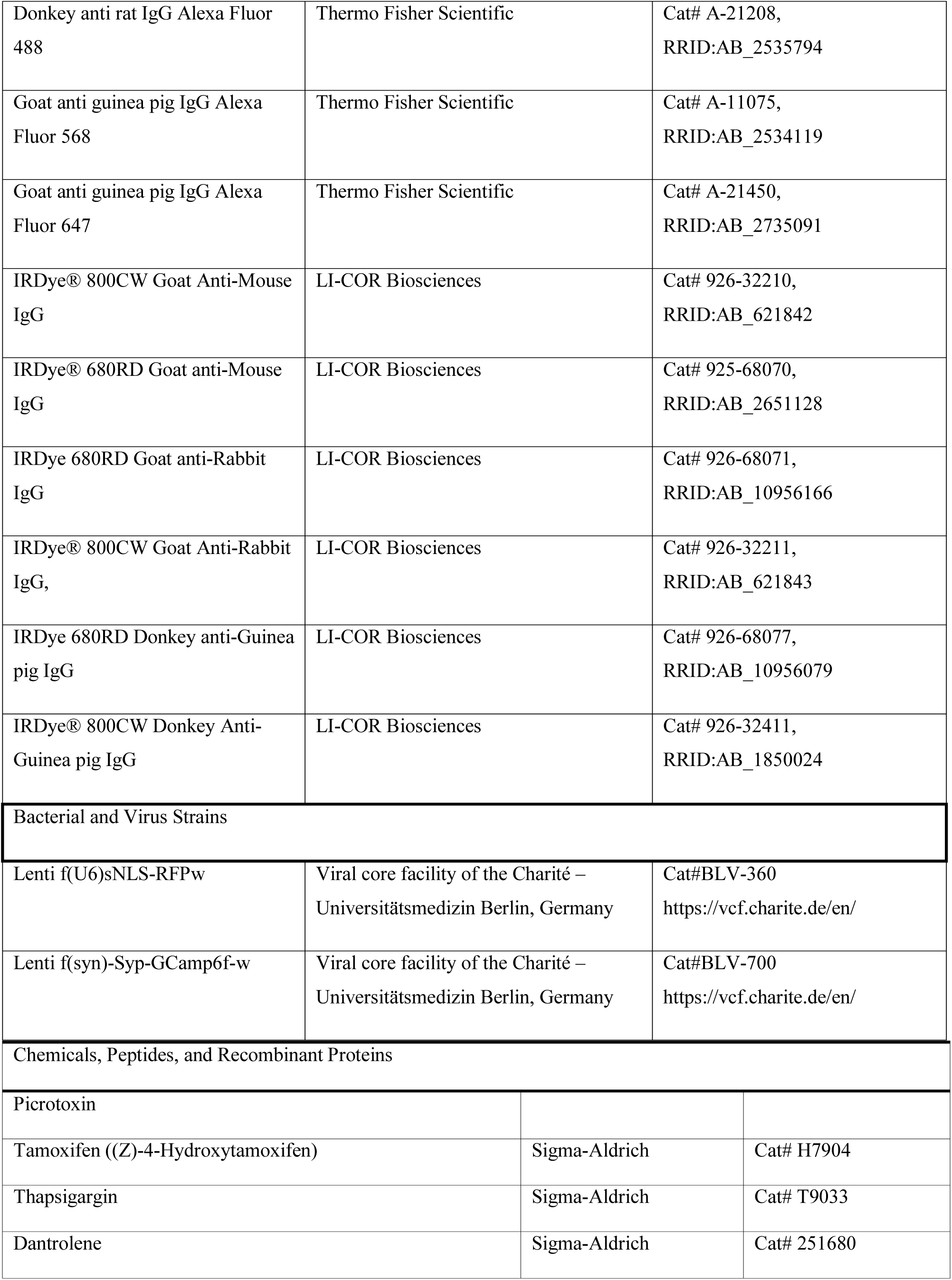
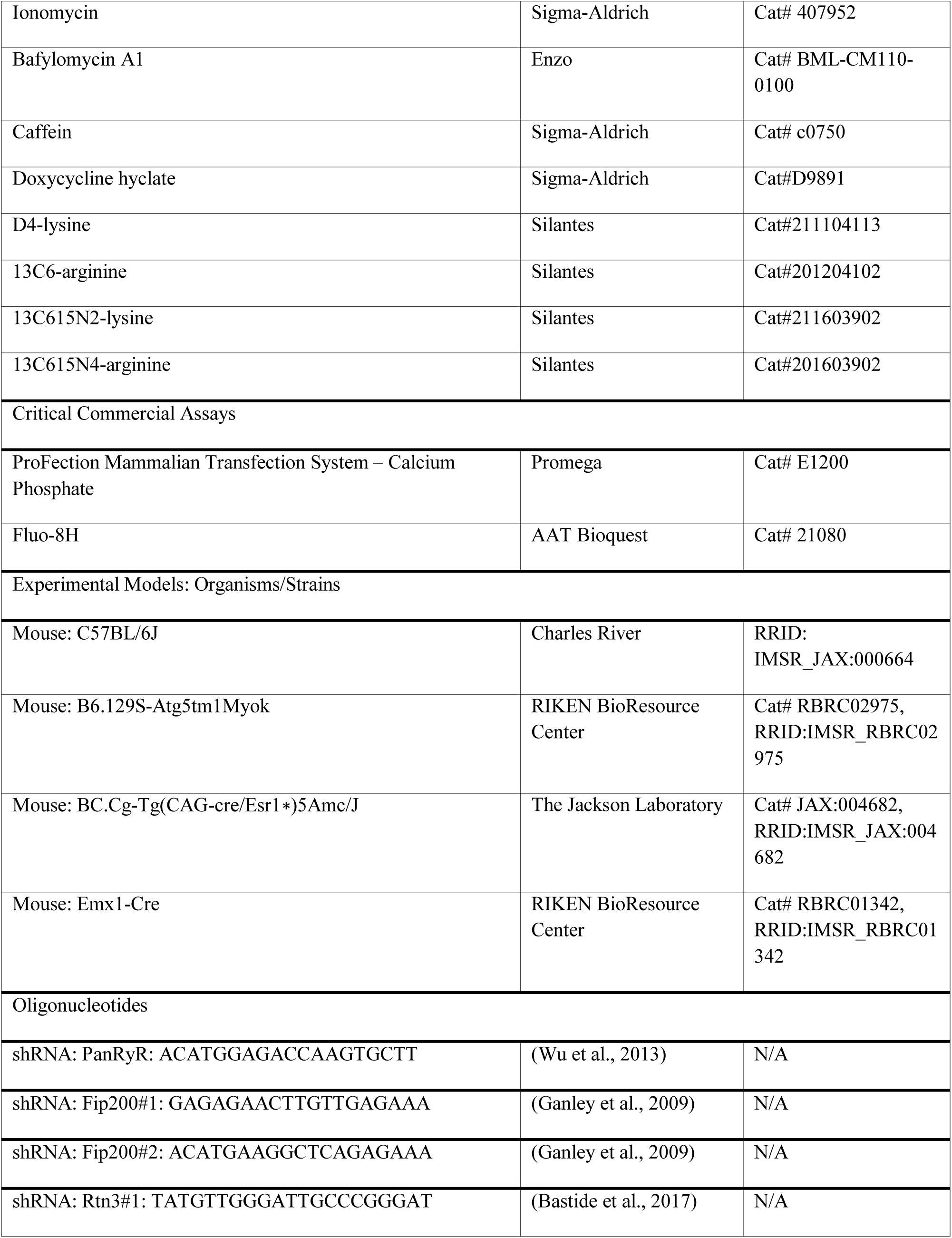
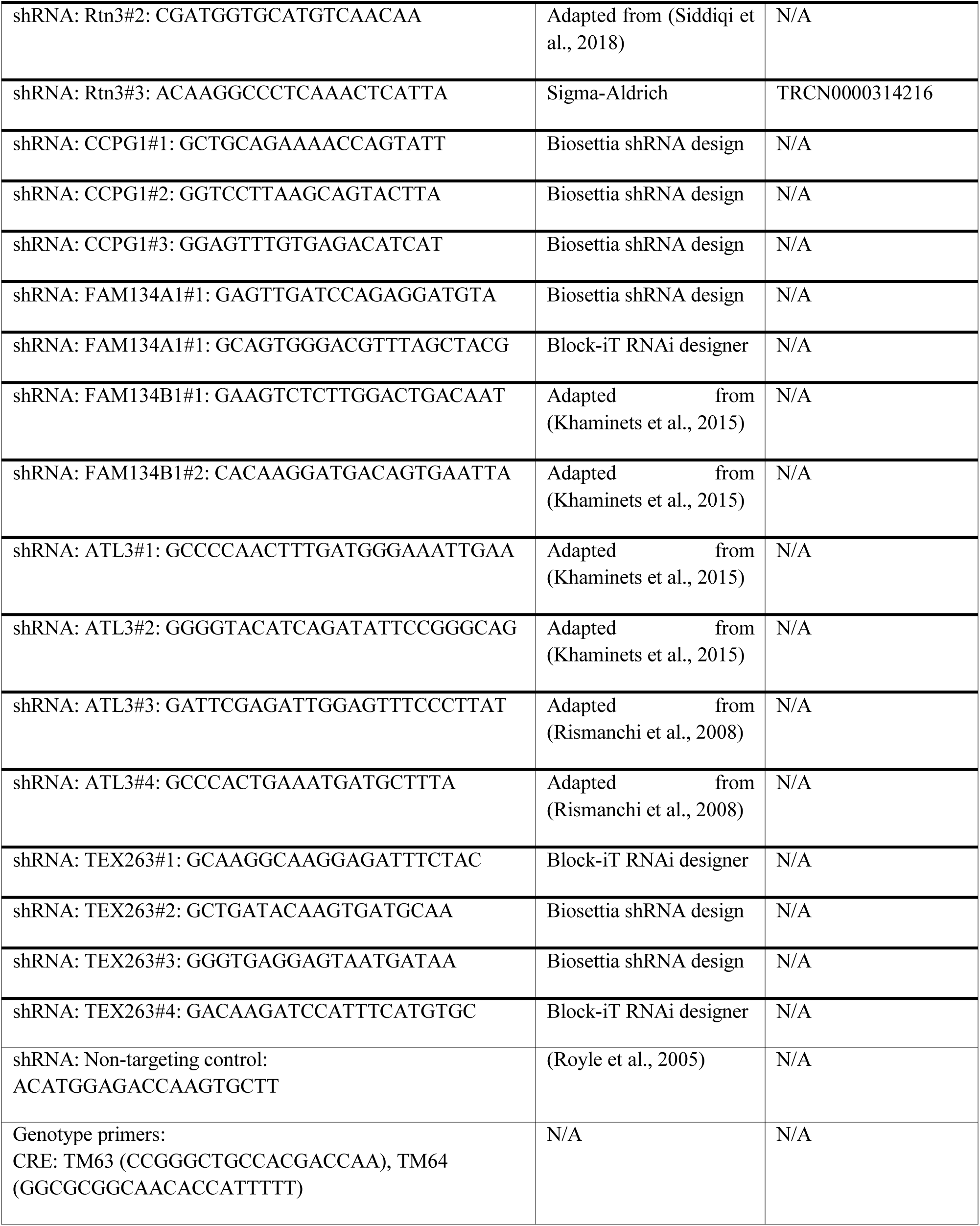
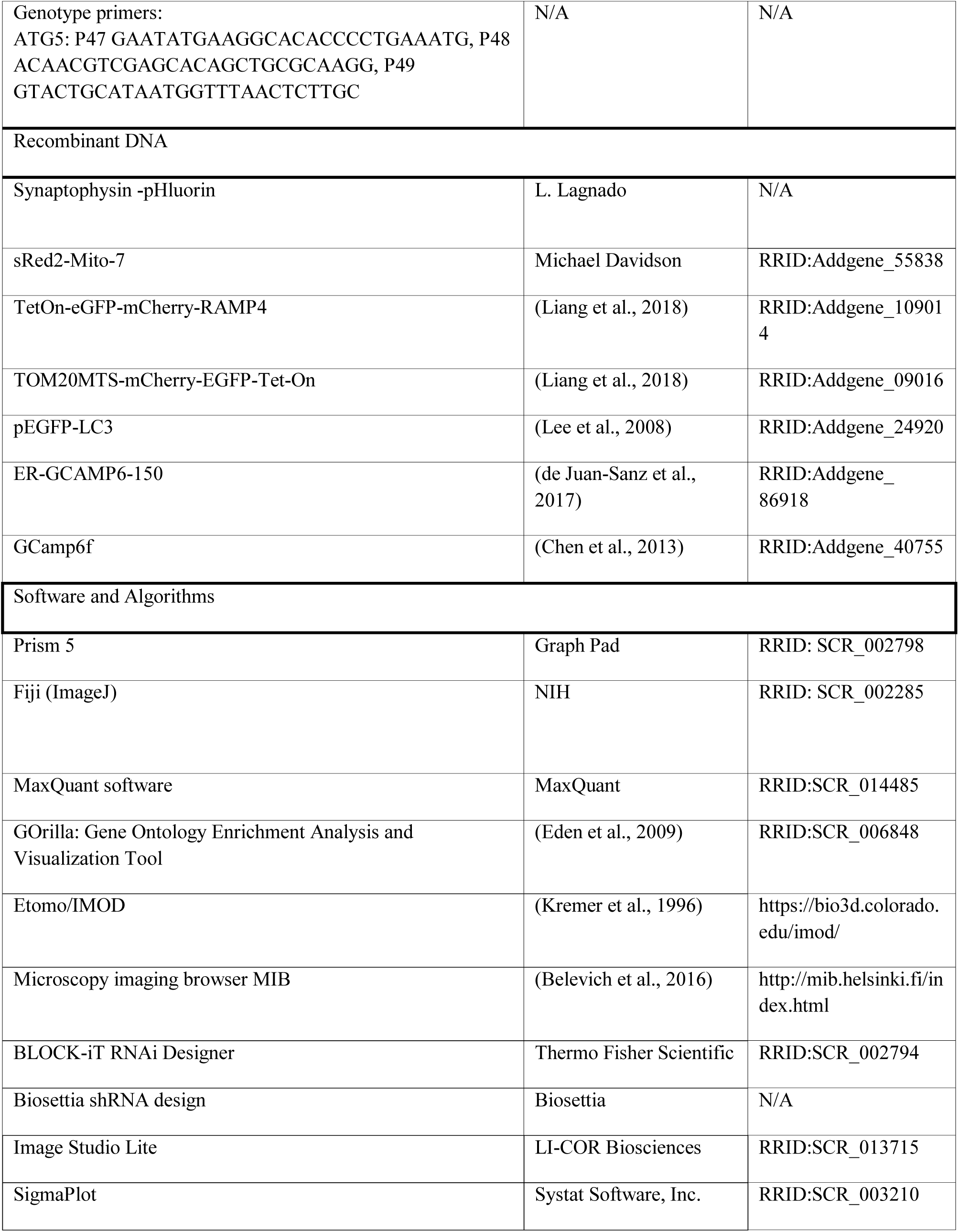
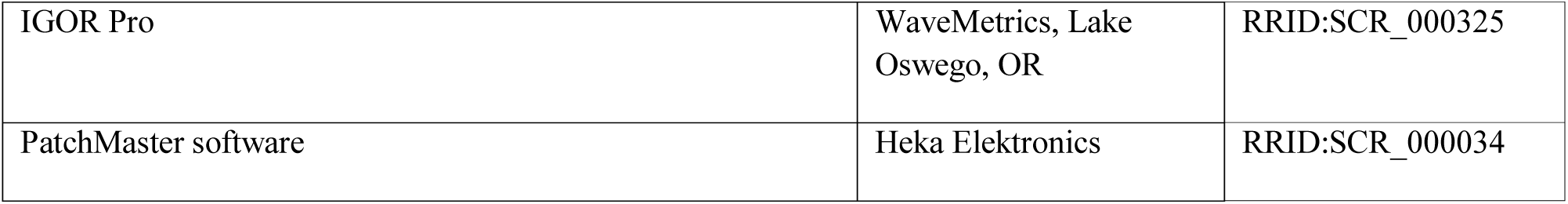

