## Supplement for "Neuronal autophagy regulates presynaptic neurotransmission by controlling the axonal endoplasmic reticulum"

### Inventory of supplementary items

#### Supplementary Figures

Supplementary figure S1 (related to Figure 1) | Elevated basal synaptic transmission and decreased mf-LTP in ATG5-cKO mice

Supplementary figure S2 (related to Figures 2 and 3) | Accumlation of proteins in ATG5- iKO neurons

Supplementary figure S3 (related to Figure 4) | Loss of autophagy leads to axonal ER accumulation but does not affect Golgi, lysosomes or mitochondria

Supplementary figure S4 (related to Figures 5 and 6) | Loss of ATG5 does not affect mitochondrial acidification, axonal lipid levels or ER integrity

Supplementary figure S5 (related to Figure 6) | ATG5 KO brains have increased RYR levels and inhibiting RYR function rescues elevated neurotransmission in ATG5-iKO neurons

**Legend to Supplementary Table S1** SILAC-based quantitative proteomic analyses shows nearly 2,000 neuronal proteins that do not show significant changes in turnover after ATG5 depletion (first tab) and a selection of proteins that exhibited a reduced degradation rate (second tab).

### Supplementary figures

S1 (Related to Figure 1)

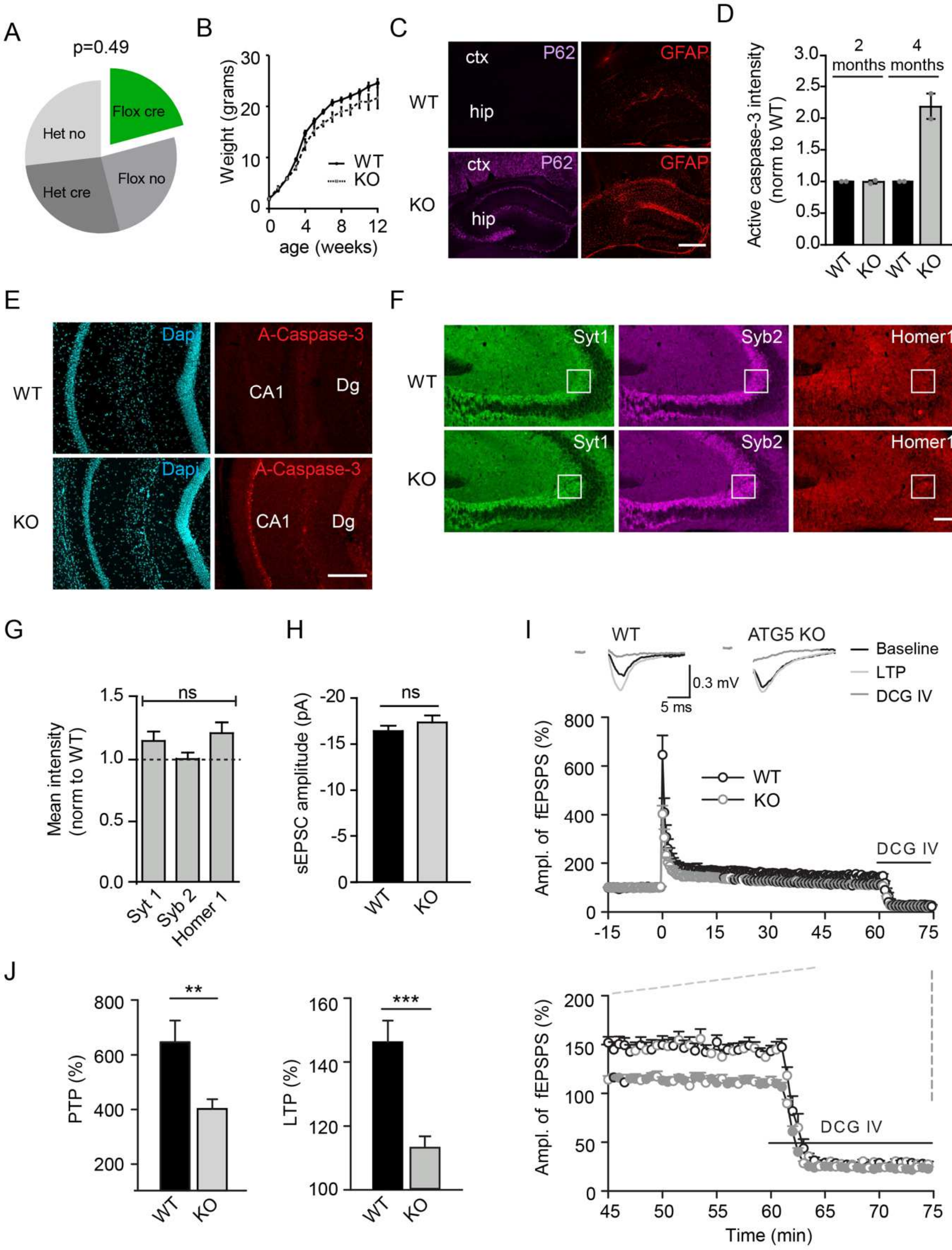

**Supplementary figure S1 (related to Figure 1) | Elevated basal synaptic transmission and decreased mf-LTP in ATG5-cKO mice**

(A) Birth ratios of ATG5$^{flox/flox}$ x ATG5$^{flox/-}$;EMX1-*Cre* matings with chi-squared analysis. n=235 mice.

(B) Growth curves of ATG5-cKO (n=4-44) mice and their littermate controls (n=16-103).

(C) GFAP and p62 immunostaining in 6-7 week old control and ATG5-cKO brain slices. Ctx=cortex, hip=hippocampus. Scale bar, 400 µm.

(D,E) Active caspase-3 immunostaining in young (<2months) versus "old" (4 months) control and ATG5-cKO brain slices. (D) Quantification of active caspase-3 intensity in the CA1 area. Slices taken from 2 mice. The mean value for the controls are set to 1, and the mean value for the KO is expressed relative to this. Values of single mice are plotted as individual points in the graph. (E) An example of active caspase-3 immunostaining in "old" (4 months) mice. Dg=dentate gyrus. Scale bar, 200 µm.

(F,G) Immunostaining in 6-7 week old control and ATG5-cKO brain slices. (F) Images of hippocampal sections depicting typical levels of synaptotagmin 1 (Syt1), synaptobrevin 2 (Syb2) and homer 1 immunoreactivities. White boxes indicate area taken for quantification (G) Expression levels of Syt1, Syb2 and Homer1 are not significantly different in ATG5-cKO compared to control littermates. The mean value for the controls was set to 1, and the mean value for the KO is expressed relative to this. Slices taken from 3 mice; one-sample t-test. Scale bar, 100 µm.

(H) Summary result of averaged sEPSC amplitude. n=14 (WT) or 17 (KO) slices from 4 animals; Mann–Whitney test. (I) Decreased post-tetanic and long-term potentiation induced by 5xHFS at mossy fiber-CA3 synapses in Atg5-cKO (n=8 slices, 6 mice) as compared to control mice (n=6 slices, 5 mice). Representative MF-fEPSP traces (above) are collected before (black), 45-60 min after LTP induction (gray) and after application of the agonist of type II metabotropic glutamate receptors DCG IV (2 μM; dark gray). Only responses inhibited by 70-80% and more were assumed to be elicited by mossy fiber synapses. The mean amplitudes of fEPSPs recorded between -15 to 0 min was taken as 100%. (J) PTP and LTP levels measured immediately after induction (WT; 645.6 ± 79.4 and Atg5-cKO; 402.4 ± 34.9) and between 45–60 min (WT; 146.2 ± 6.7 and Atg5-cKO; 113.2 ± 3.6), respectively, show reduced facilitation in Atg5-cKO mice; unpaired t-test.

All data represent mean ± SEM, ns: not significant, ∗∗p < 0.01; ∗∗∗p < 0.001.

S2 (Related to Figure 2 and 3)

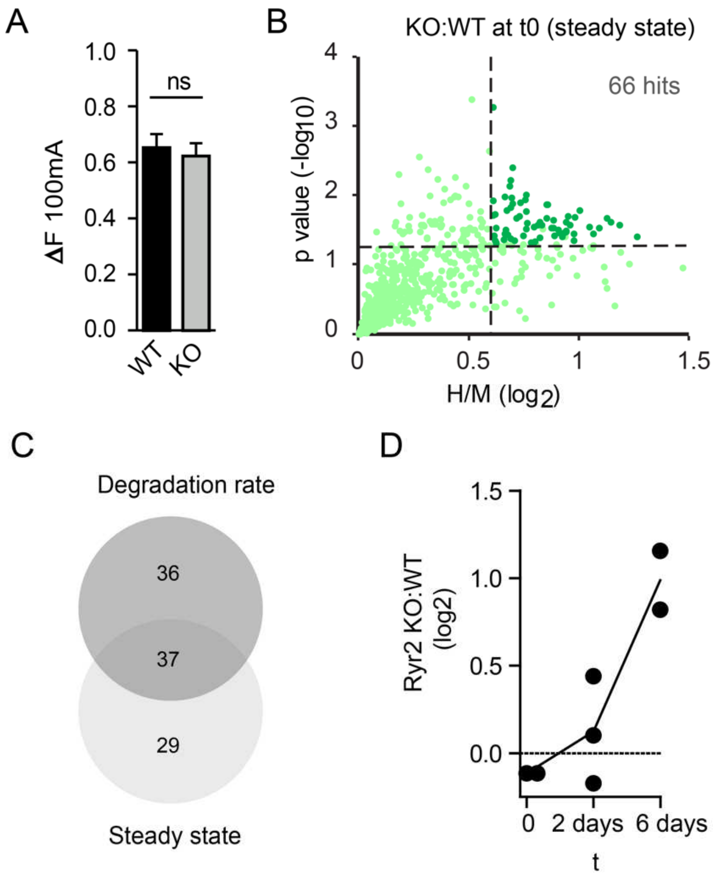


**Supplementary figure S2 (related to Figures 2 and 3) | Accumlation of proteins in ATG5-iKO neurons**

(A) Maximal fluorescent peak of synaptophysin–pHluorin–expressing neurons at 100 mA (Fmax, 60AP). n=28-30 cells, 4 independent experiments; unpaired t-test. Data represent mean ± SEM, ns: not significant. See also Figure 2E.

(B) Comparisons of H/M (KO/WT) ratios obtained for proteins of neuron cultures that are lysed at t=0 (DIV14, no replacement with unlabeled medium). Instead of differences in turnover (see

Figure 3D for pulsed SILAC) this graph rather shows the accumulation of proteins in the ATG5-iKO neurons (defined as KO/WT>1.5 and $p<0.05$, dotted lines).

(C) Overlap of protein hits found in steady state SILAC vs pulsed SILAC

(D) Ryanodine Receptor 2 (RyR2) was not detected in all SILAC conditions (and therefore not one of 1753 detected proteins) but plotting of H/M (WT/KO) ratios of single conditions (black dots) shows a clear trend of slower RyR2 degradation in ATG5-iKO neurons.

S3 (Related to Figure 4)

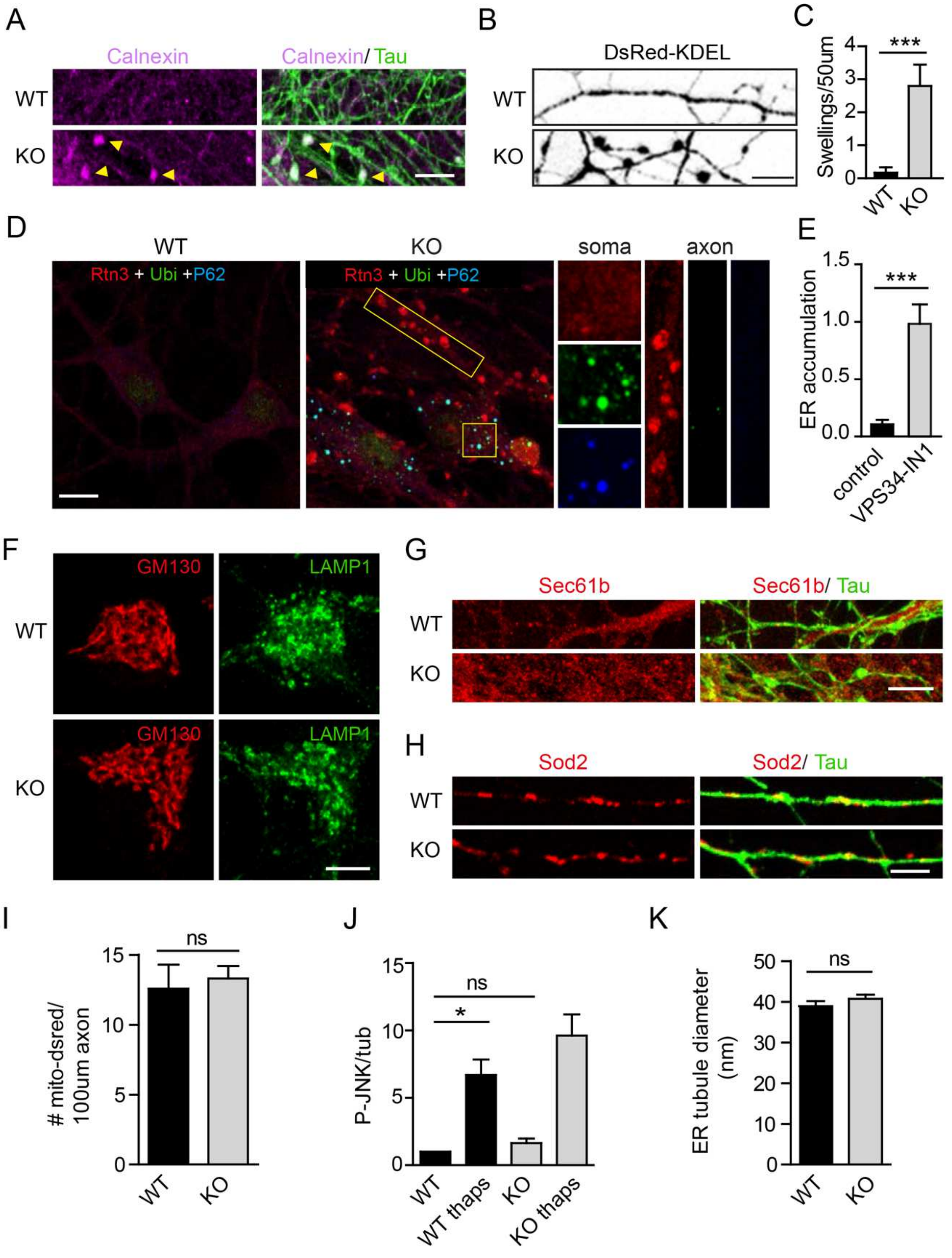

**Supplementary figure S3 (related to Figure 4) | Loss of autophagy leads to axonal ER accumulation but does not affect Golgi, lysosomes or mitochondria**

(A) Representative confocal images of WT and ATG5-iKO hippocampal neurons immunostained for ER marker Calnexin and axonal marker Tau. Yellow arrows indicate ER accumulations in KO axons.

(B,C) WT and ATG5-iKO hippocampal neurons transfected with DsRed-KDEL. (B) Representative confocal images of DsRed-KDEL positive axons. (C) Quantification of amount of KDEL positive swellings. n=12 (WT) or n=10 (KO) images, 1 experiment; unpaired t-test.

(D) Hippocampal neurons stained for Rtn3, Ubiquitin (Ubi) and P62. Enlarged regions (yellow boxes) show P62 and ubiquitin accumulations in soma and Rtn3 accumulation in axons.

(E) Quantification of Rtn3 in control and VPS34-IN1 (1 μM, 24 hours) treated neurons, expressed as axonal area (Tau positive, not shown) covered with Rtn3 accumulations. n=16-17 images, 1 experiment; unpaired t-test.

(F) Representative confocal images of WT and ATG5-iKO hippocampal neurons immunostained for Golgi marker GM130 and lysosome marker LAMP1.

(G) Representative confocal images of WT and ATG5-iKO hippocampal neurons immunostained for rough ER marker Sec61.

(H-I) No apparent differences between WT and ATG5-iKO neurons in immunostainings and transfection of mitochondrial markers (H) Number of axonal mitochondria quantified in WT and AT5 KO neurons transfected with dsred-mito, n=3 experiments, 26 images per condition; paired t-test

(J) Quantification of JNK phosphorylation (p-JNK) in WT and ATG5-iKO neuron lysates, with and without thapsigargin treatment to induce ER stress (thaps, 1 μM). Conditions are compared to WT, values for WT were set to 1. n=4 experiments; one-sample t-test. See also Figure 4F.

(K) ER tubule diameter measured in electron micrographs of WT and ATG5-iKO hippocampal neurons. n=61 tubules from 13 images (WT) or n=62 tubules from 8 images (KO); unpaired t-test. Scale bars, 5 µm. All data represent mean ± SEM, ns: not significant, $*p < 0.05$; $***p < 0.001$.

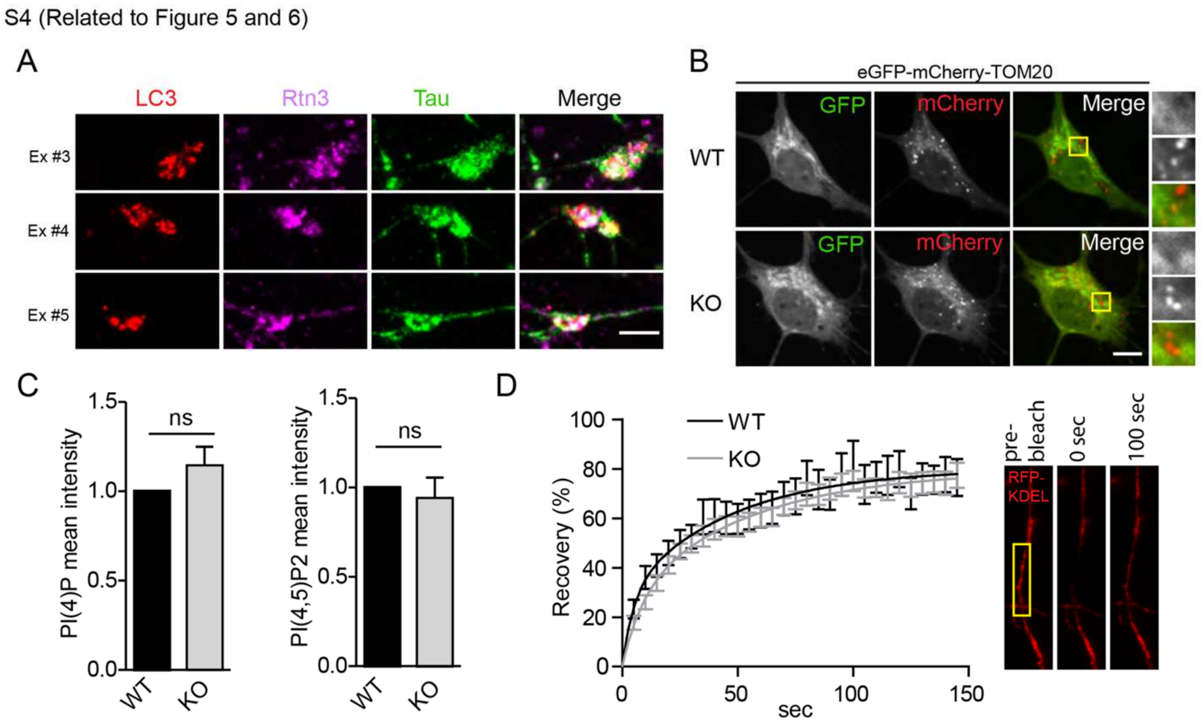


**Supplementary figure S4 (related to Figures 5 and 6) | Loss of ATG5 does not affect mitochondrial acidification, axonal lipid levels or ER integrity**

(A) Additional examples of VPS34 inhibitor washout experiment described in Figure 5G. Scale bar, 10 µm

(B) Representative confocal images of hippocampal neurons transfected with eGFP-mCherry-TOM20. eGFP is quenched as a result of low pH, causing a switch from GFP+/mCherry+ to GFP−/mCherry+ during lysosomal degradation of mitochondria. Yellow boxes indicate magnifications shown on the right. Scale bar, 5 µm. See Figure 5J for quantification.

(C) Quantification of PI(4)P and PI(4,5)P2 immunostainings in WT and KO synapses (marked by VAMP2 immunostaining). The mean values for the WT are set to 1. n=3 (PI(4)P) or n=4 (PI(4,5)P2) independent experiments; one-sample t-test.

(D) Fluorescent recovery plots showing the rates of DsRed-KDEL recovery in axons of control neurons and ATG5-iKO neurons. Fluorescent intensity was normalized to intensity before bleaching. n=15 (WT) or 21 (KO) axons, the black and grey curves are the fit of a nonlinear regression model to the experimental data. Images on the right show a time series example, the photobleached area is indicated with a yellow box.

All data represent mean ± SEM, ns: not significant.

S5 (Related to Figure 6)

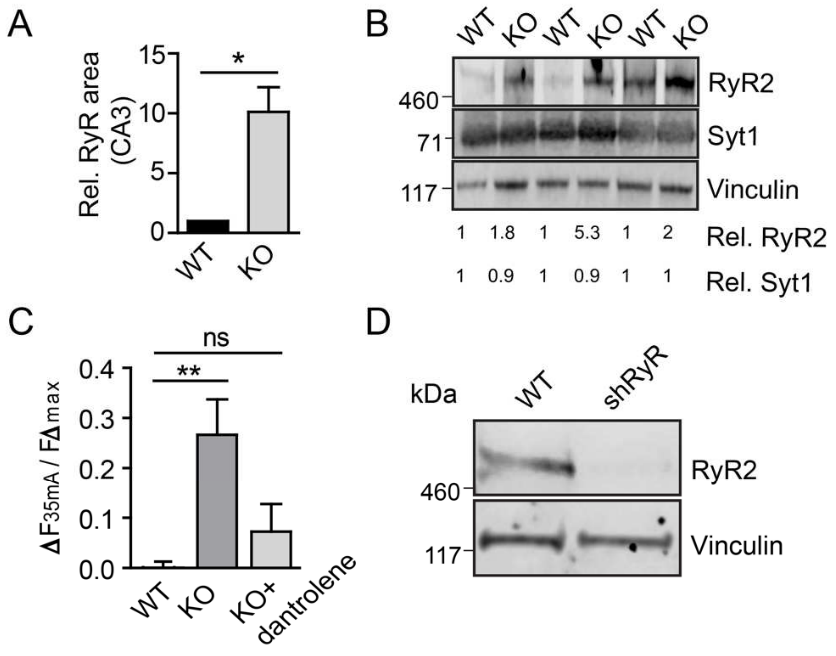


E

| shRNA | Axonal ER accumulation |
|---|---|
| FIP200 | yes |
| Reticulon-3 | no |
| FAM134A | no |
| FAM134B | no |
| CCPG1 | no |
| TEX264 | no |
| Atlastin-3 | no |

**Supplementary figure S5 (related to Figure 6) | ATG5 KO brains have increased RYR levels and inhibiting RYR function rescues elevated neurotransmission in ATG5-iKO neurons**

(A) Ryanodine Receptor immunoreactivity in hippocampal CA3 area of WT and ATG5-cKO mice. n=3; one-sample t-test. See also Figure 6E.

(B) Immunoblots of WT and ATG5-cKO brain lysates showing increased Ryanodine Receptor 2 (RyR2) levels and no difference in synaptotagmin 1 (Syt1) levels. Density measurements of indicated proteins (normalized to housekeeping protein vinculin) are indicated below the blots.

(C) Detection of exocytosis using synaptophysin-pHluorin in WT and ATG5-iKO hippocampal neurons. Graph showing mean normalized peak fluorescence upon a 35mA stimulation. Dantrolene (10 µM), a Ryanodine Receptor inhibitor, rescues increased responses in ATG5-iKO neurons. Values per cell are normalized to the corresponding maximal fluorescent peak at 100 mA (Fmax). n=18-22 cells, 3 experiments; one-way ANOVA with Tukey's post-test.

(D) Immunoblot of lysates from neurons infected with scrambled or (pan)RYR-shRNA virus, probed for RYR2 and vinculin antibodies.

(E) Lentiviral –mediated knockdown (KD) screen for the indicated ER-phagy adaptors. None of the adaptor KD conditions (each consisting of 2-4 different shRNA sequences) lead to axonal ER increases measured by calnexin or Rtn3 immunostainings.

All data represent mean ± SEM, ns: not significant, $*p < 0.05$; $**p < 0.01$.
